# Defining Inflammatory Cell States in Rheumatoid Arthritis Joint Synovial Tissues by Integrating Single-cell Transcriptomics and Mass Cytometry

**DOI:** 10.1101/351130

**Authors:** Fan Zhang, Kevin Wei, Kamil Slowikowski, Chamith Y. Fonseka, Deepak A. Rao, Stephen Kelly, Susan M. Goodman, Darren Tabechian, Laura B. Hughes, Karen Salomon-Escoto, Gerald F. M. Watts, William Apruzzese, David J. Lieb, David L. Boyle, Arthur M. Mandelin, Accelerating Medicines Partnership: RA Phase 1, AMP RA/SLE, Brendan F. Boyce, Edward DiCarlo, Ellen M. Gravallese, Peter K. Gregersen, Larry Moreland, Gary S. Firestein, Nir Hacohen, Chad Nusbaum, James A. Lederer, Harris Perlman, Costantino Pitzalis, Andrew Filer, V. Michael Holers, Vivian P. Bykerk, Laura T. Donlin, Jennifer H. Anolik, Michael B. Brenner, Soumya Raychaudhuri

## Abstract

To define the cell populations in rheumatoid arthritis (RA) driving joint inflammation, we applied single-cell RNA-seq (scRNA-seq), mass cytometry, bulk RNA-seq, and flow cytometry to sorted T cells, B cells, monocytes, and fibroblasts from 51 synovial tissue RA and osteoarthritis (OA) patient samples. Utilizing an integrated computational strategy based on canonical correlation analysis to 5,452 scRNA-seq profiles, we identified 18 unique cell populations. Combining mass cytometry and transcriptomics together revealed cell states expanded in RA synovia: *THY1*^+^*HLA*^*high*^ sublining fibroblasts (OR=33.8), *IL1B*^+^ pro-inflammatory monocytes (OR=7.8), *CD11c*^+^*T-bet*^+^ autoimmune-associated B cells (OR=5.7), and *PD-1*^+^Tph/Tfh (OR=3.0). We also defined CD8^+^ T cell subsets characterized by *GZMK*^+^, *GZMB*^+^, and *GNLY*^+^ expression. Using bulk and single-cell data, we mapped inflammatory mediators to source cell populations, for example attributing *IL6* production to *THY1*^+^*HLA*^*high*^ fibroblasts and naïve B cells, and *IL1B* to pro-inflammatory monocytes. These populations are potentially key mediators of RA pathogenesis.

---

Rheumatoid arthritis (RA) is an autoimmune disease affecting up to 1% of the population where a complex interplay between many different cell types drives chronic inflammation in the synovium of the joint tissue^1–3^. This inflammation leads to joint destruction, disability and shortened life span^4^. Defining key cellular subsets and their activation states in RA has been a longstanding key step to defining new therapeutic targets. CD4^+^ T cell subsets^5,6^, B cells^7^, monocytes^8,9^, and fibroblasts^10–12^ have established relevance to RA pathogenesis. A global portrait of RA-relevant cell subsets using single cell technologies across a large sample collection tissues from inflamed joints is a critical resource for advancing therapeutics.

Application of transcriptomic and cellular profiling technologies to whole synovial tissue has already identified promising specific cellular populations associated with RA^3,13–15^. However, most studies have focused on a pre-selected cell type, surveyed whole tissues rather than disaggregated cells, or used only one technology. Latest advances in single-cell technologies offer an opportunity to identify disease-associated cell subsets in human tissues at high resolution in an unbiased fashion^16–19^. These technologies have already indicated roles for T peripheral helper (Tph) cells^20^ and HLA-DR+CD27^−^ cytotoxic T cells^21^ in RA pathogenesis. Separately, scRNA-seq has defined myeloid cell heterogeneity in human blood^22^ and identified a distinct subset of PDPN^+^CD34^−^THY1^+^ (THY1, also known as CD90) fibroblasts enriched in RA synovial tissue^16,23^.

To generate high-dimensional multi-modal single-cell data from synovial tissue samples, we developed a robust tissue analytical pipeline^24^ in the Accelerating Medicines Partnership (AMP) RA/SLE consortium. We collected and disaggregated tissue samples from patients with RA and OA, and then subjected constituent cells to scRNA-seq, sorted-population bulk RNA-seq, mass cytometry, and flow cytometry. We developed a robust computational strategy based on canonical correlation analysis (CCA) to integrate multi-modal transcriptomic and proteomic profiles at a single cell level. A unified analysis of single cells across data modalities can precisely define contributions of specific cell subsets to pathways relevant to RA and chronic inflammation.

## RESULTS

### Generation of parallel mass cytometric and transcriptomic data from synovial tissue

In phase 1 of AMP-RA/SLE, we recruited 36 RA patients meeting 1987 ACR classification criteria and 15 OA control patients from 10 clinical sites over 16 months (**Supplemental Table 1**) and obtained synovial tissues from ultrasound guided synovial biopsies or joint replacements (**Methods**). All tissue samples included had with synovial lining documented by histology (**Fig. 1a**). Synovial tissue disaggregation yielded many viable cells (362,190 cells per tissue, S.E.M 7,687 cells) for downstream analyses. Applying a previously validated strategy for synovial cell sorting^24^ (**Fig. 1a**), we separated cells into B cells (CD45^+^CD3^−^CD19^+^), T cells (CD45^+^CD3^+^), monocytes (CD45^+^CD14^+^), and stromal fibroblasts (CD45^−^PDPN^+^) (**Supplemental Fig. 1a**). We applied bulk RNA-seq to all four sorted subsets from the 51 samples. For a subset of samples with sufficient cell yield (Methods), we measured single-cell protein expression using a 34-marker mass cytometry panel (n=26, **Supplemental Table 2**), and single-cell RNA expression in sorted populations (n=21, **Fig. 1b**).

**Fig. 1.**
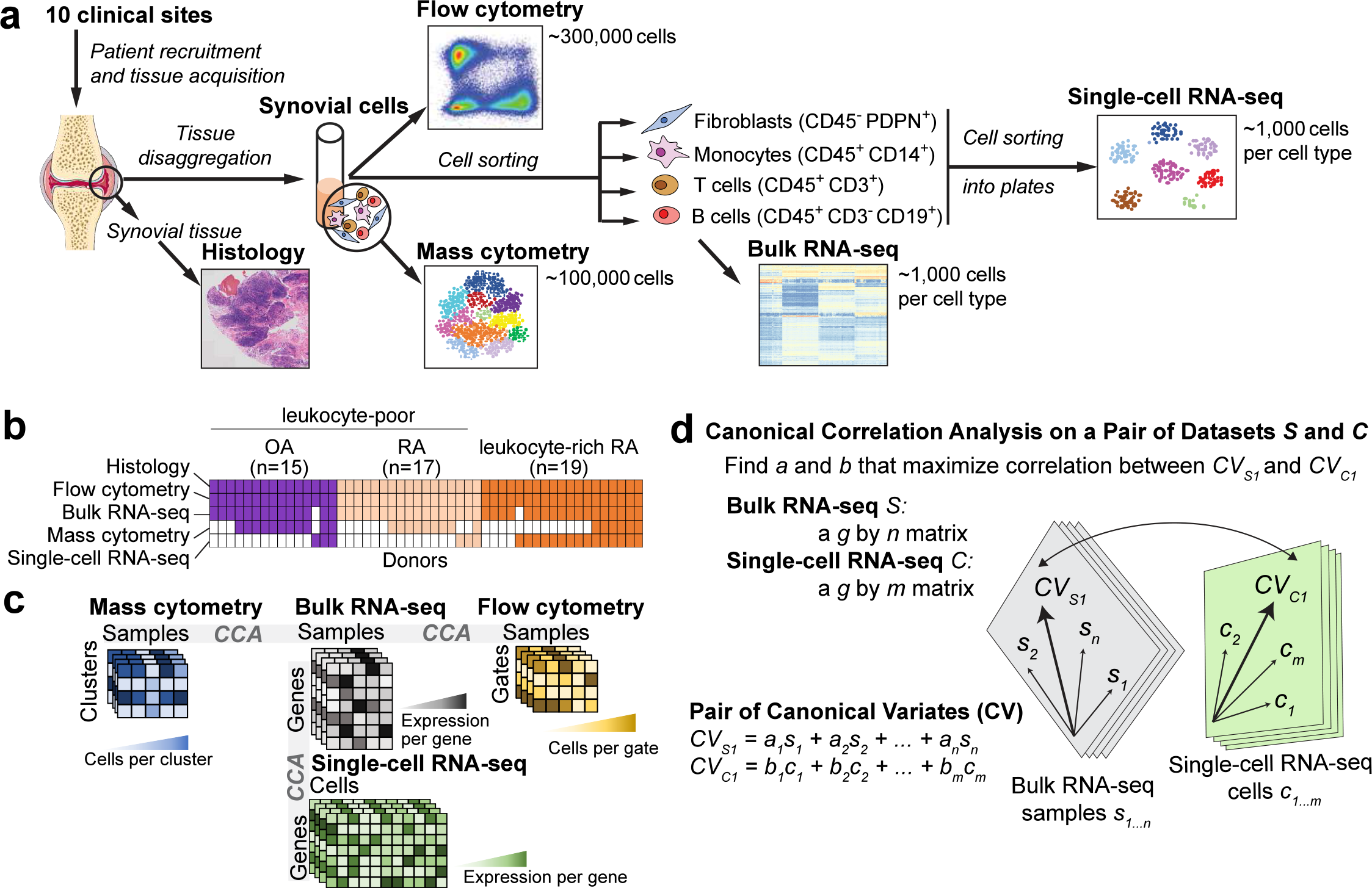
Overview of synovial tissue workflow and pairwise analysis of high-dimensional data. **a.** We acquired synovial tissue, disaggregated the cells, sorted them into four gates representing fibroblasts (CD45^−^PDPN^+^), monocytes (CD45^+^CD14^+^), T cells (CD45^+^CD3^+^), and B cells (CD45^+^CD3^−^CD19^+^). We profiled these cells with mass cytometry, flow cytometry, sorted low-input bulk RNA-seq, and single-cell RNA-seq. **b.** Presence and absence of five different data types for each tissue sample, **c.** Schematic of each dataset and the shared dimensions used to analyze each of the 3 pairs of datasets with canonical correlation analysis (CCA), **d.** CCA finds a common mapping for two datasets. For bulk and single-cell RNA-seq, we first find a common set of g genes present in both datasets. Each bulk sample *s_j_* gets a coefficient *a_i_* and each cell *c_i_*, gets a coefficient *b_i_*. The linear combination of all samples *s_1…n_* arranges bulk genes along the canonical variate *CV_S1_* and the linear combination of all cells *c_1…m_* arranges single-cell genes along *CV_C1_*. CCA finds the coefficients *a_1…n_* and *b_1…m_* that arrange the genes from the two datasets in such a way that the correlation betwen the genes is maximized. After CCA finds the first pair of canonical variates, the next pair is computed on the residuals, and so on.

### Summary of computational data integration strategy to define cell populations

To confidently define RA associated cell populations, we used bulk RNA-seq data as the reference point for our study (**Fig. 1c**). Bulk RNA-seq data were available for almost all of the samples, had the highest dimensionality and least sparsity, and were the least sensitive to technical artifacts.

We used CCA to integrate bulk RNA-seq data with the three other datasets (**Fig. 1c**). Integrating scRNA-seq with bulk RNA-seq data ensures robust discovery of individual cellular populations. Here, we used CCA to find linear combinations of bulk RNA-seq samples and scRNA-seq cells (Fig. 1d) to create gene expression profiles that were maximally correlated. These linear combinations captured sources of shared variation between the two datasets and allowed us to identify individual cellular populations that drive variation in the bulk RNA-seq data. We clustered scRNA-seq data by using the most correlated canonical variates for each cell to compute a nearest neighbor network, and then identified clusters with a community detection algorithm (**Methods, Supplemental Fig. 2a**).

We identified clusters of cells in mass cytometry data using density-based clustering^25^. To define the genes that best correspond to the mass cytometry clusters, we integrated bulk RNA-seq with mass cytometry using CCA. In this analysis, CCA identifies linear combinations of genes and mass cytometry cluster proportions so that correlation across individual samples is maximized. These canonical variates offer a way to visualize genes and mass cytometry clusters together and define genes possibly specific for individual clusters. We then integrated mass cytometry clusters with identified scRNA-seq clusters to define the relationship between them (**Methods**). We also associated bulk gene expression in each sample with proportions of cells in different flow cytometry gate by integrating bulk RNA-seq with flow cytometry data using CCA.

### Disease association test of cellular populations

We tested whether abundances of individual populations were altered in RA case samples compared to controls using two ways. First, we assessed whether marker genes (AUC>0.7, 20 < n < 100) of each scRNA-seq derived cluster was differentially expressed concordantly in bulk RNA-seq samples. Second, we applied MASC^21^, a single cell association testing framework, to identify mass cytometry clusters associated with disease (**Methods**).

### Synovial lymphocyte and monocyte infiltration distinguishes leukocyte-rich RA synovia

Histology of RA synovial tissues revealed heterogeneous tissue composition with variable lymphocyte and monocyte infiltration (**Fig. 2a,b**); in contrast OA tissues had minimal lymphocytic infiltration (**Fig. 2a**). This expected heterogeneity reflects variable disease activity among RA patients which results in differences in tissue immune cell infiltration^26^. Consequently, we employed a data-driven approach to separate samples based on the degrees of lymphocyte and monocyte infiltration of tissues measured by flow cytometry (**Supplemental Fig. 1b,c**). We calculated a multivariate normal distribution of these parameters based on OA samples as a reference, and then for each RA sample calculated the Mahalanobis distance from OA^27^. We defined the maximum OA value (4.5) as a threshold to separate all leukocyte-rich RA samples from leukocyte-poor samples (**Methods, Supplemental Fig. 1d**). We defined 19 leukocyte-rich RA and 17 leukocyte-poor RA samples in our cohort. Whereas leukocyte-rich RA tissues had marked infiltration of synovial T cells and B cells (**Fig. 2c**), leukocyte-poor RA tissues had a similar cellular composition of leukocytes and stromal fibroblasts as OA (**Fig. 2c**). Synovial monocytes were similar between RA and OA (**Fig. 2c**).

**Fig. 2.**
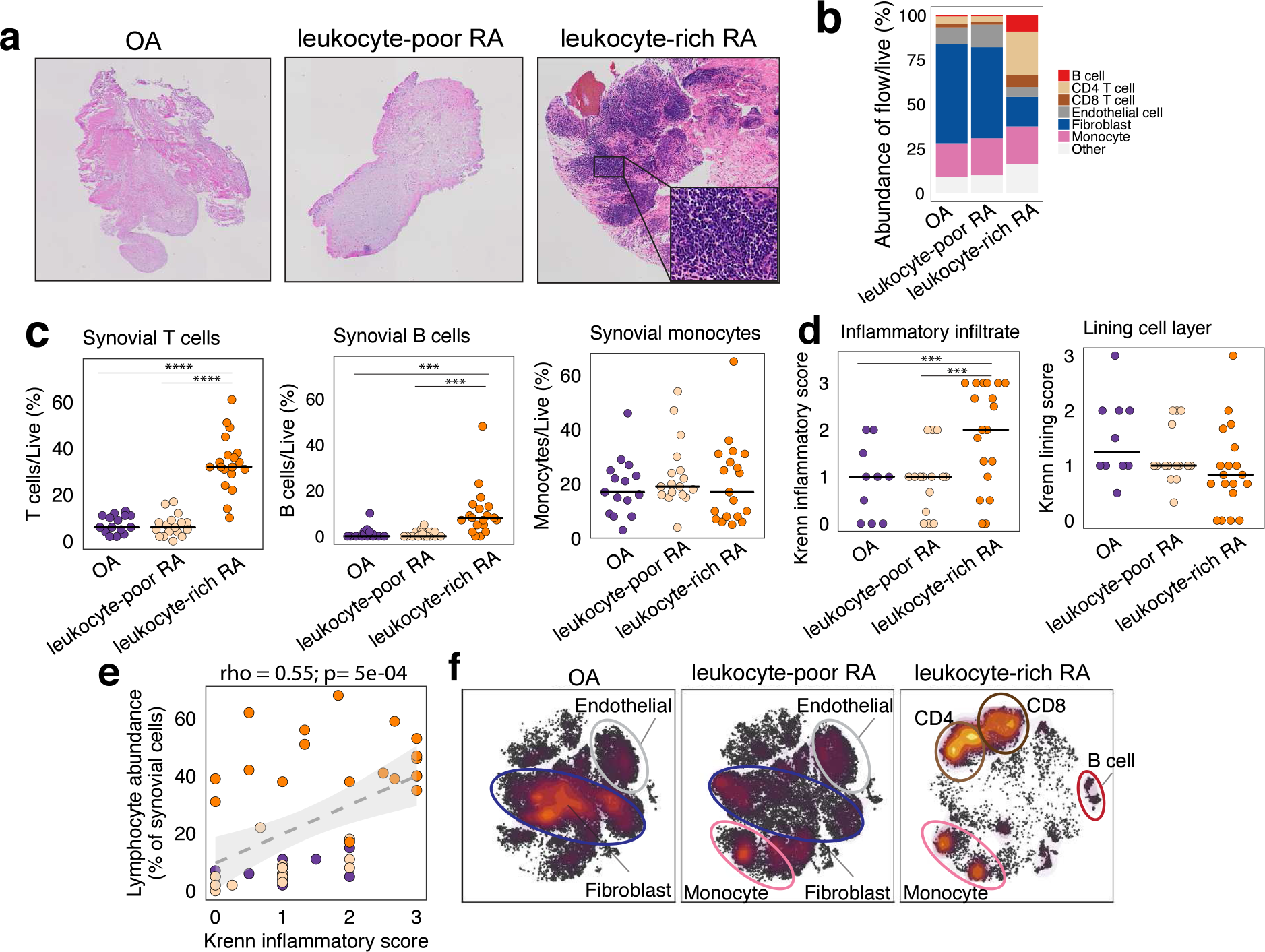
Distinct cellular composition in synovial tissue from OA, leukocyte-poor RA, and leukocyte-rich RA patients, **a.** Histological assessment of synovial tissue derived from OA, leukocyte-poor RA, and leukocyte-rich RA. **b.** Cellular composition of major synovial cell types by flow cytometry, **c.** Synovial T cells, B cells, and monocytes by flow cytometry in samples from OA (n=15), leukocyte-poor RA (n=17), and leukocyte-rich RA (n=19). Leukocyte-rich RA tissues were significantly higher infiltrated in synovial T cells (Student’s t-test p=4e-9, t-value=8.92, *61=22.27*) and B cells (Student’s t-test p=1e-3, t-value=3.50, df=20.56) compared to OA. Statisitcal significance levels: ****p ≤ 1e-4, ***p ≤ le-3. **d.** Quantitative histologic inflammatory scoring of both sublining cell layer and lining layer. Leukocyte-rich RA samples (n=19) exhibited higher (Student’s t-test p=1 e-3, t-value=3.21, df=30.66) Krenn inflammation scores than leukocyte-poor RA(n=15) and OA tissues (n=10) samples, **e.** Spearman correlation (rho = 0.55, p=5e-04) between lymphocytic infiltration assessed by cytometry with histologic inflammation score (n=44). **f.** tSNE visualization of synovial cell types in OA, leukocyte-poor RA, and leukocyte-rich RA by mass cytometry density plot.

Mass cytometry in 26 synovial tissues was consistent with flow cytometric and histologic analyses. We observed marked differences in synovial cellular composition between leukocyte-rich RA, leukocyte-poor RA, and OA. Stromal fibroblasts and endothelial cells constituted most synovial cells in OA and leukocyte-poor RA and are otherwise characterized by expansion of monocytes with few lymphocytes (**Fig. 2f**, **Supplemental Fig. 3**). In stark contrast, leukocyte-rich RA tissues constituted predominantly of CD4 T, CD8 T, and B cells (**Fig. 2f**).

To validate whether our classification indicated inflammation, we assessed tissue histology and assigned a Krenn inflammation score^28^. We observed that leukocyte-rich RA samples exhibited significantly higher score than leukocyte-poor RA and OA (**Fig. 2d**). In contrast, synovial lining membrane hyperplasia was not significantly different between leukocyte-rich RA, leukocyte-poor RA, and OA controls. We observed significant correlation between synovial lymphocyte infiltration and histologic inflammation score (*t*-test p=5e-04; Spearman’s rho = 0.55, **Fig. 2e**), suggesting consistent classification between cytometric and histologic assessments.

### Single-cell RNA-seq analysis reveals distinct cell subpopulations

Next, we analyzed 5,265 scRNA-seq profiles passing stringent quality control, including 1,142 B cells, 1,844 fibroblasts, 750 monocytes, and 1,529 T cells (**Methods**). We used canonical variates (from bulk RNA-seq integration) to define clusters that were independent of donor and sequencing batch effects (**Fig. 3a-b**, **Supplemental Fig. 2b,c**). In contrast, conventional PCA-based clustering led to clusters that were confounded by batch effects (**Supplemental Fig. 2d,e**). We selected marker genes for scRNA-seq clusters by comparing cells within it to cells outside it and applied the following criteria: 1) percent of non-zero expressing cells > 60%; 2) AUC score > 0.7; and 3) FC > 2 (**Supplemental Table 4**). CCA-based clustering identified 18 clusters (4 fibroblast clusters, 4 monocyte clusters, 6 T cell clusters, and 4 B cell clusters) from 21 donors (**Fig. 3a**, interactive form at https://immunogenomics.io/amp/). The distribution of these distinct clusters varies between donors, suggesting heterogeneity in immune and stromal cell subsets across patients (**Fig. 3b**). We show typical markers for cells in a t-Distributed Stochastic Neighbor Embedding (tSNE^29^) into two-dimensional space (**Fig. 3c-f**). Here we briefly summarize these populations.

**Fig. 3.**
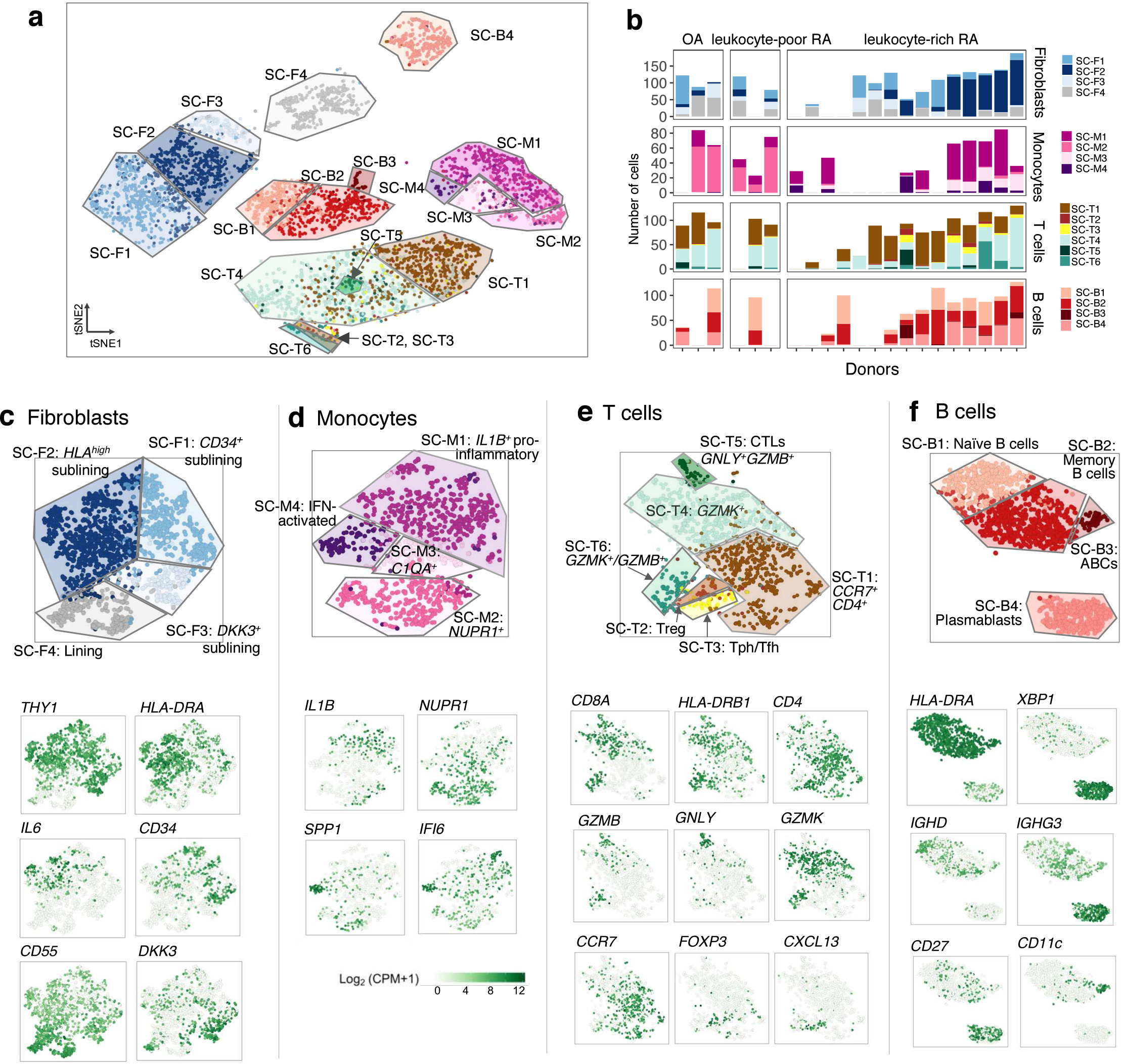
High-dimensional transcriptomic single-cell RNA-seq clustering reveals distinct cell type subpopulations, **a.** Clustering of 5,265 cells from all cell types reveals 18 distinct cell type subpopulations, **b.** Distribution of identified cell clusters reveals heterogeneity of individual donors, **c.** Distinct synovial fibroblast subsets including three types of *THY1^+^*sublining fibroblasts (SC-F1, SC-F2, and SC-F3) and *CD55^+^* lining fibroblasts (SC-F4). **d.** Distinct monocyte subsets including two activated cell states of *IL1B\** pro-inflammatory (SC-M1) and IFN-activated (SC-M4) monocytes, **e.** Heterogeneity in synovial T cells: *CD4^+^* subsets: SC-T1, SC-T2, SC-T3, and CDS^+^ subsets: SC-T4, SC-T5, and SC-T6. **f.** Distinct B cells subsets including *HLA^+^* (SC-B1, SC-B2, and SC-B3) and plasmablasts (SC-B4). The cluster colors in **c-f** are consistent with the colors in the clustering of all the cells (**a**).

Within stromal fibroblasts, we identified four putative cell subpopulations (**Fig. 3c**). The *CD55^+^* (SC-F4) cluster represented lining fibroblasts and were the most different from the other fibroblast clusters^16,23^. The other three fibroblast clusters were *CD34*^+^ sublining fibroblasts (SC-F1), *HLA*^*high*^ sublining fibroblasts (SC-F2), and *DKK3*^+^ sublining fibroblasts (SC-F3). In monocytes (**Fig. 3d**), we identified *IL1B*^+^ pro-inflammatory monocytes (SC-M1), *NUPR1*^+^ monocytes (SC-M2), *C1QA*^+^ monocytes (SC-M3), and IFN-activated monocytes (SC-M4). In T cells (**Fig. 3e**), we identified three CD4^+^ clusters: *CCR7*^+^*CD4*^+^ T cells (SC-T1), *FOXP3*^+^ Tregs (SC-T2), and *PD-1*^+^Tph/Tfh (SC-T3). We also found three CD8^+^ clusters: *GZMK*^+^ T cells (SC-T4), *GNLY*^+^*GZMB*^+^ cytotoxic lymphocytes (CTLs) (SC-T5), and *GZMK*^+^/*GZMB*^+^ T cells (SC-T6). Within B cells (**Fig. 3f**), we identified four cell clusters, including naive *IGHD*^+^*CD27*^−^ (SC-B1) and *IGHG3*^+^*CD27*^−^ memory B cells (SC-B2). Intriguingly, we identified an autoimmune-associated B cell (ABC) cluster (SC-B3) with high expression of *ITGAX* (CD11c)^30,31^. We also identified a plasmablast cluster (SC-B4) with high expression of *IgG* genes *and XBP1*, a transcription factor critical for plasma cell differentiation^32^.

### Distinct synovial fibroblasts defined by cytokine activation and MHC II expression

In synovial fibroblasts, differential single cell gene expression suggested that CD55^+^ fibroblasts (SC-F4) were the most transcriptionally distinct subset from the three sublining *THY1^+^* clusters SC-F1, SC-F2, and SC-F3, indicating that anatomical localization contributes to synovial fibroblast diversity^16,23^ (**Fig. 4a**). Consistent with the role of synovial fibroblasts in matrix remodeling, the three sublining fibroblasts, *CD34^+^* fibroblasts (SC-F1), *HLA^high^* fibroblasts (SC-F2), and *DKK3^+^* fibroblasts (SC-F3) share gene expression in pathways related to extracellular matrix constituents by gene set enrichment analysis (GSEA) (**Fig. 4a,b**). *HLA^high^* sublining fibroblasts (SC-F2) are enriched with genes related to MHC class II presentation (*HLA-DRA* and *HLA-DRB1*) and the interferon gamma-mediated signaling pathway (*IFI30*) (**Fig. 4a,b**), suggesting upregulation of MHC class II in response to interferon-gamma signaling in these cells. We identified a novel sublining fibroblast subtype (SC-F3) that is characterized by high expression of *DKK3, CADM1* and *COL8A2*.

**Fig. 4.**
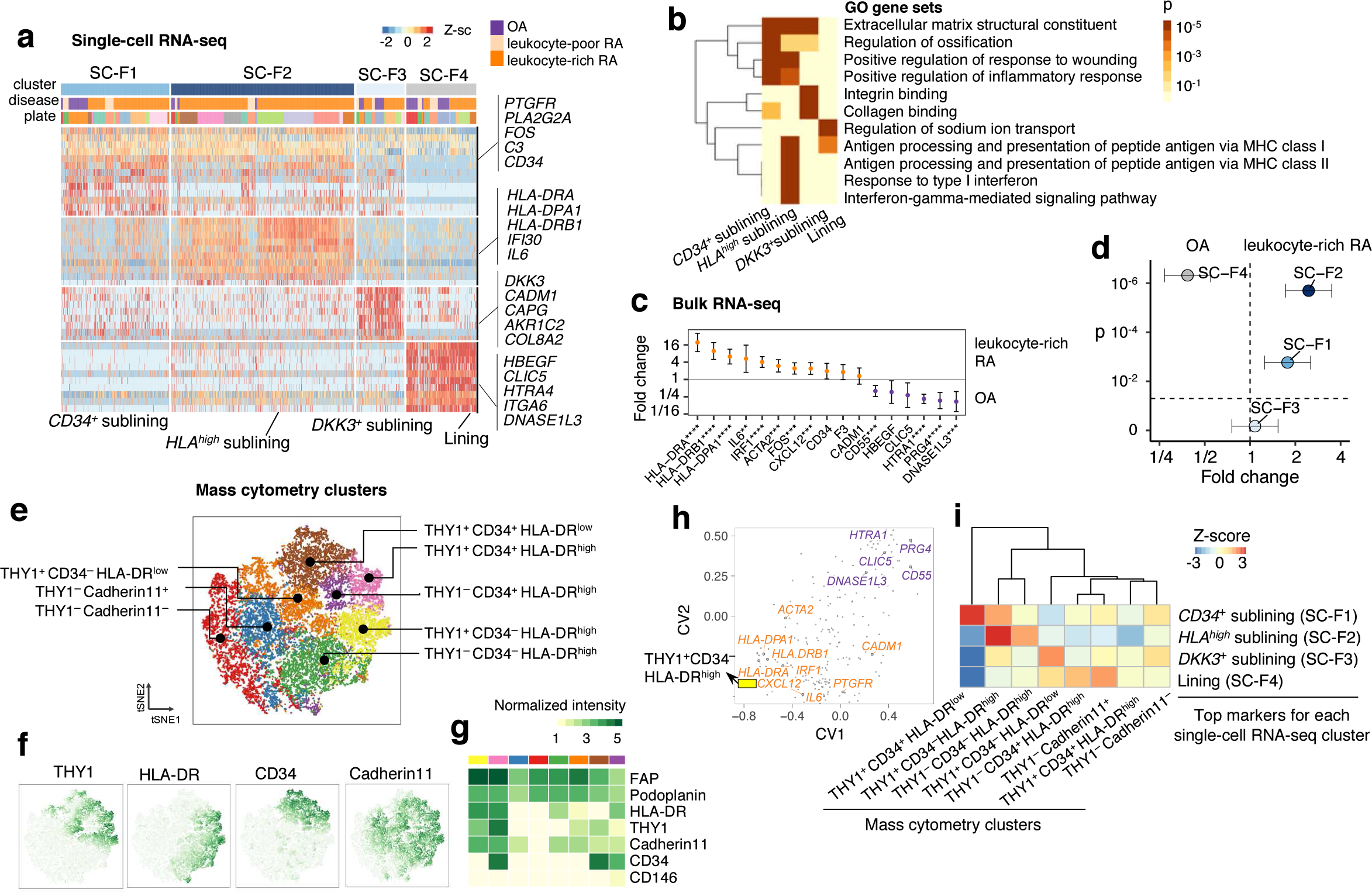
Distinct synovial fibroblasts defined by cytokine activation and MHC II expression, **a.** Single-cell RNA-seq analysis identified three sublining fibroblasts, *CD34^+^* (SC-F1), *HLA^high^* (SC-F2), and *DKK3*^+^ (SC-F3) and one lining subset (SC-F4). **b.** Pathway enrichment analysis indicates the potential pathways for each cluster, **c.** Differential analysis on leukocyte-rich RA (n=16) with OA (n=12) by bulk RNA-seq fibroblast samples revealed genes that upregulated and downregulated in leukocyte-rich RA. Effect size with 95% confidence intervals are given. Significantly differential expressed (****p ≤ le-4, ***p ≤ 1e-3, **p ≤ 1e-2) genes are highlighted, **d.** By querying the leukocyte-rich RA (n=16) and OA (n=12) fibroblast bulk RNA-seq samples, single-cell RNA-seq cluster *HLA^+^* (SC-F2) and *CD34^+^* (SC-F1) fibroblasts are significantly upregulated (two-tailed Student’s *t*-test p=2e-6, t-value=6.2, df = 23.91 and p=2e-3, t-value = 3.20, df = 25.41, respectively) in leukocyte-rich RA, while lining cells (SC-F4) are enriched (two-tailed Student’s *t*-test p=5e-7, t-value=−5.31, df =21.97) in OA samples, **e.** Mass cytometry analysis revealed eight distinct populations, **f-g**. Normalized intensity of distinct protein markers are shown in tSNE visualization and heatmap. h. Integration of identified mass cytometry clusters with bulk RNA-seq using CCA. First two canonical variates (CV) separated genes that upregulated in leukocyte-rich RA from genes that depleted in leukocyte-rich RA. *HLA^high^* genes are highly associated with THY1^+^CD34^−^HLA-DR^high^ by mass cytometry, **i.** Integration of mass cytometry clusters with single-cell RNA-seq clusters using the top markers (AUC > 0.7) for each single-cell RNA-seq cluster based on the top 10 canonical variates in the CCA space. We computed the spearman correlation between each pair of single-cell RNA-seq cluster and mass cytometry cluster in the CCA space and performed permutation test 10^4^ times. Z-score is calculated based on permutation p-value. We observed *HLA^high^* sublining fibroblasts are strongly correlated with THY1 ^+^CD34^−^HLA-DR^high^ fibroblasts by mass cytometry.

To identify fibroblast populations expanded in leukocyte-rich RA synovia, we first examined expression of genes associated with each fibroblast subsets from bulk-sorted fibroblasts (CD45^−^PDPN^+^) from RA and OA patients. Expression of genes associated with *HLA^high^* fibroblasts (*HLA-DRA, IRF1, ACTA2*, and *CXCL12*, *t*-test p<1e-3) were upregulated in leukocyte-rich RA (n=16) compared to OA (n=12) by bulk RNA-seq (**Fig. 4c**), suggesting expansion of SC-F2. Genes associated with SC-F4 lining fibroblasts (*PRG4, CD55, HTRA1*, and *DNASE1L3*, *t*-test p<1e-3) were significantly decreased in leukocyte-rich RA (**Fig. 4c**). Next, we used the most differentially expressed genes (AUC>0.7) in each fibroblast subset to query transcriptomic profiles of bulk-sorted fibroblasts from leukocyte-rich RA and OA synovia. *HLA^high^* sublining fibroblasts (SC-F2) and *CD34^+^* sublining fibroblasts (SC-F1) were significantly expanded in RA synovia compared to OA (*t*-test p=2.5e-6 and p=2.1e-3, respectively), while *CD55^+^* lining fibroblasts (SC-F4) were relatively decreased in leukocyte-rich RA (*t*-test p=5.0e-7) (**Fig. 4d**).

We then queried the proteomic expression to validate these four fibroblast populations. Analysis of CD45^−^PDPN^+^ cells identified eight putative cell clusters based on the differential expression pattern of THY1, HLA-DR, CD34, and Cadherin11 (**Fig. 4e-g**) that were not confounded by obvious batch effects (**Supplemental Fig. 4a**). Integration of mass cytometry clusters with bulk RNA-seq using CCA showed that the *IL6, CXCL12*, and *HLA* gene expression is highly associated with frequency of THY1^+^CD34^−^HLA-DR^high^ fibroblasts, suggesting an active cytokine-producing state (**Fig. 4h**). In contrast, the expression of lining fibroblast genes *PRG4* and *CD55* separated in the CCA space with a gradient, indicating relative decreased number of lining fibroblasts in leukocyte-rich synovium (**Fig. 4h**). We then integrated each scRNA-seq subset based on the most unique genes (AUC>0.7) with identified the corresponding mass cytometry clusters and determined the statistical significance (z-score) of this association by explicit permutation (**Fig. 4i**, **Methods**). We consistently observed that *HLA^high^* sublining fibroblasts (SC-F2) are strongly associated (z-score=2.8) with THY1^+^CD34^−^HLA-DR^high^ fibroblasts, and *CD34^+^* sublining fibroblasts (SC-F1) are strongly correlated (z-score=2.7) with THY1^+^CD34^+^HLA-DR^low^ fibroblasts (**Fig. 4h**, **Table 1**) indicating that these populations correspond to each other.

**Table 1.**
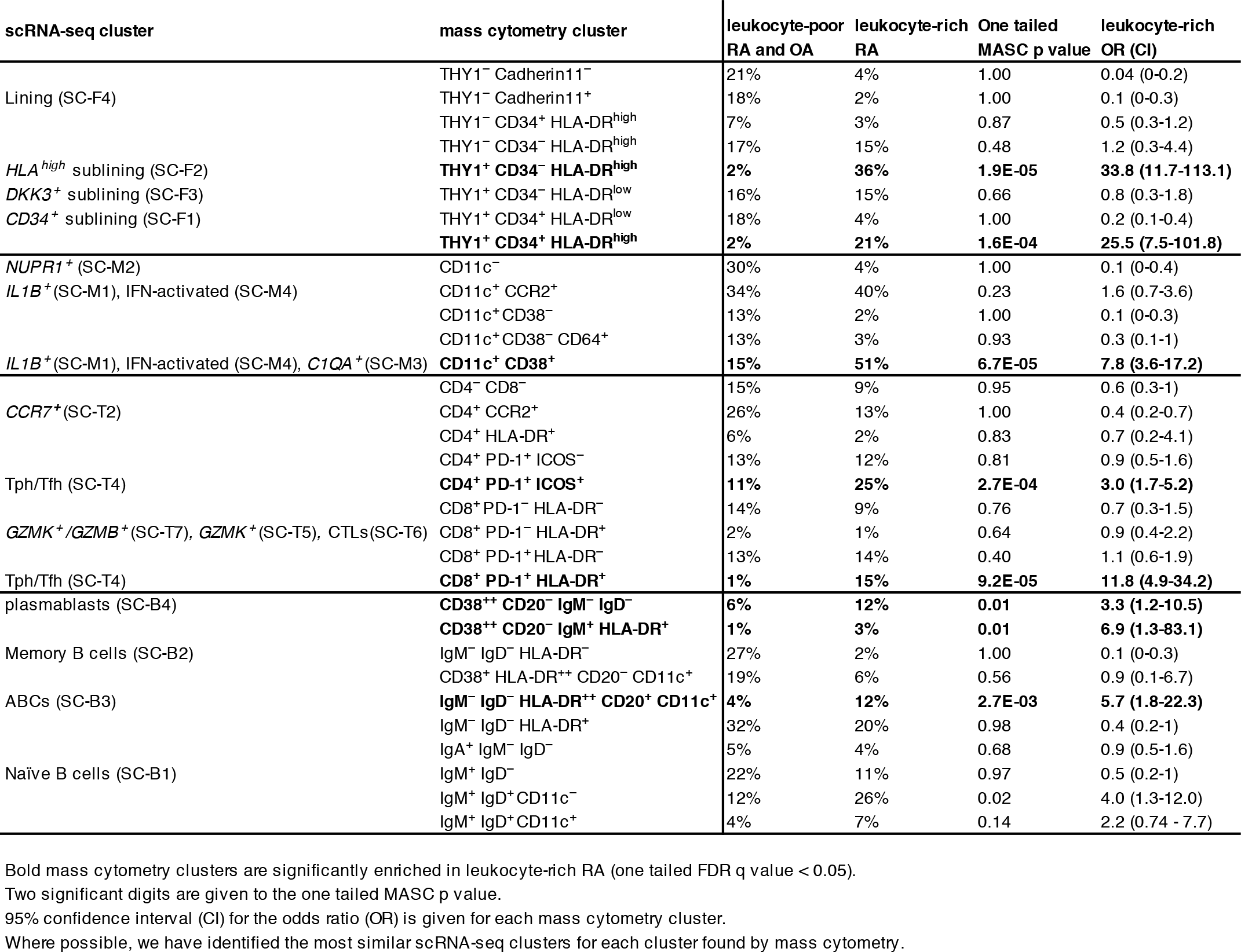
Connection between cell populations determined by mass cytometry and scRNA-seq clusters and disease associations.

Consistent with the differential expression analyses, we found that THY1^+^CD34^−^HLA-DR^high^ cells are dramatically overabundant in leukocyte-rich RA compared to leukocyte-poor RA and OA controls (36% versus 2% of fibroblasts, MASC OR = 33.8 (95% CI: 11.7-113.1), one tailed MASC p=1.9e-05) (**Table 1**).

### Unique activation states define heterogeneity among synovial monocytes

With scRNA-seq, we defined four transcriptionally distinct monocyte subsets: *IL1B^+^* pro-inflammatory monocytes (SC-M1), *NUPR1*^+^ monocytes (SC-M2), *C1QA^+^* monocytes (SC-M3) and IFN-activated monocytes (SC-M4) (**Fig. 5a**). GSEA demonstrated that monocyte LPS response was associated with SC-M1 (44.8% of total monocytes) (**Fig. 5b**), suggesting it represents a phenotype similar to IL-1- or TLR-activated proinflammatory monocytes. Using Gene Ontology gene sets, we observed that SC-M4 monocytes were highly enriched in the type I interferon signaling and the interferon-gamma mediated pathway (**Supplemental Fig. 5a**), including increased expression of *IFITM3^22^* and *IFI6* (**Fig. 5a**). The phenotypes of the monocytes from SC-M2 and SC-M3 clusters do not align well with known activation states, possibly indicating a more homeostatic role in the synovium.

**Fig. 5.**
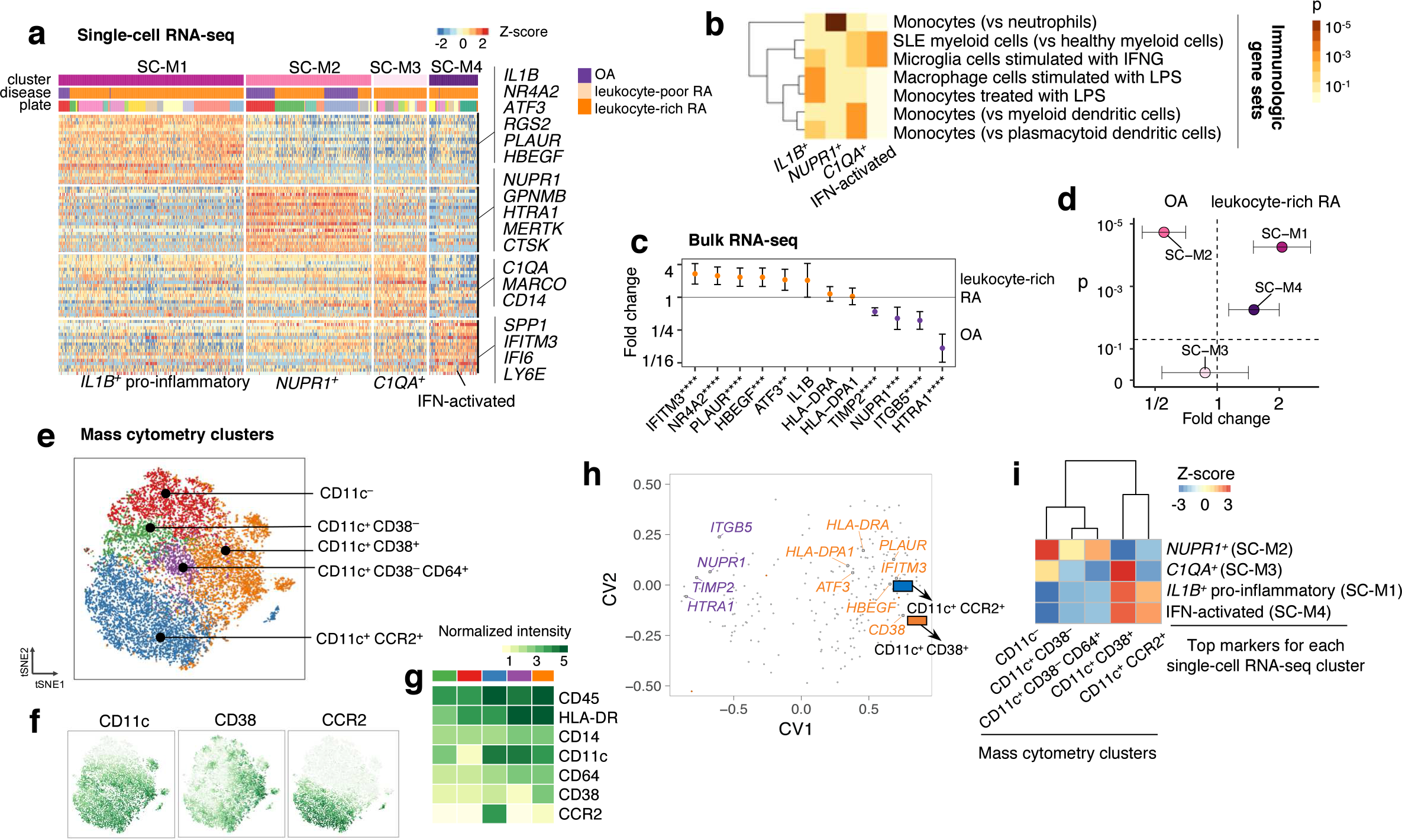
Unique activation states define synovial monocytes heterogeneity, **a.** Single-cell RNA-seq analysis identified four subsets: *IL1B_*_* pro-inflammatory monocytes (SC-M1), *NUPR1^+^* monocytes (SC-M2) with a mixture of RA and OA cells, *C1QA^+^* (SC-M3), and IFN-activated monocytes (SC-M4). **b.** Pathway enrichment analysis indicates the potential pathways for each cluster. The standard names for the immunological gene sets from up to bottom are: Genes down-regulated in neutrophils versus monocytes (GSE22886); Genes down-regulated in healthy myeloid cells versus SLE myeloid cells (GSE10325); Genes down-regulated in control microglia cells versus those 24 h after stimulation with IFNG (GSE1432); Genes down-regulated in unstimulated macrophage cells versus macrophage cells stimulated with LPS (GSE14769); Genes up-regulated monocytes treated with LPS versus monocytes treated with control IgG (GSE9988); Genes up-regulated in monocytes versus myeloid dendritic cells (mDC) (GSE29618); Genes up-regulated in monocytes versus plasmacytoid dendritic cells (pDC) (GSE29618). c. Differentially expressed genes (****p ≤ 1e-4, ***p ≤ 1e-3, **p ≤ 1e-2) by bulk RNA-seq on leukocyte-rich RA samples (n=17) and OA samples (n=13). Effect size with 95% confidence intervals are given, **d.** By querying the bulk RNA-seq, we found single-cell RNA-seq cluster *IL1B\** pro-inflammatory monocytes (two-tailed Student’s *t*-test p=6e-5, t-value=4.56, df =26.33) and IFN-activated monocytes (two-tailed Student’s t-test p=6e-3, t-value=3.28, df =23.68) are upregulated in leukocyte-rich RA, while SC-M2 is depleted (two-tailed Student’s t-test p=2e-5, t-value=-5.62, df=26.81) in leukocyte-rich RA samples, **e.** Mass cytometry analysis revealed five distinct populations, **f-g.** Normalized intensity of distinct protein markers are shown in tSNE visualization and heatmap. **h.** Integration of identified mass cytometry clusters with bulk RNA-seq reveals genes that are associated with CD11c^+^CD38^+^ and CD11c^+^CCR2^+^, like *IFITM3, CD38, HBEGF, ATF3*, and *HLA^+^* genes, **i.** Integration of mass cytometry clusters and single-cell RNA-seq clusters revealed that CD11c^+^CD38^+^ and CD11c^+^CCR2^+^ by mass cytometry are significantly associated with *IL1B^+^* pro-inflammatory (SC-M1) and IFN-activated (SC-M4) monocytes.

By querying bulk RNA-seq monocyte samples from leukocyte-rich RA (n=17) and OA samples (n=13), we found that genes associated with *IL1B^+^* monocytes (SC-M1), including *NR4A2* (*t*-test p=2.2e-05), *HBEGF* (*t*-test p=1.2e-4), *PLAUR* (*t*-test p=1.5e-4) and the IFN-activated monocytes gene *IFITM3* (*t*-test p=9.3e-05) were significantly upregulated in leukocyte-rich RA samples. In contrast, marker genes associated with *NUPR1^+^* monocytes (SC-M2) were relatively depleted in leukocyte-rich RA (**Fig. 5c**). Extensive examination of the top differentially expressed genes (AUC>0.7) for each monocyte subset confirmed a significant enrichment of *IL1B^+^* monocytes (*t*-test p=6.1e-5) and IFN-activated monocytes (*t*-test p=6.2e-3) in leukocyte-rich RA synovia in contrast to a relative depletion of *NUPR1^+^* monocytes (*t*-test p=2.2e-5) (**Fig. 5d**). These data indicate that cytokine activation drives the expansion of unique monocyte populations in active RA synovia.

Mass cytometry identified five synovial CD14^+^ monocyte clusters (CD45^+^CD3^−^) (**Fig. 5e-g**) without obvious batch effects (**Supplemental Fig. 4b**). A CCA-based integration of mass cytometry and bulk RNA-seq data indicated that monocyte genes enriched in RA subsets, such as *IFITM3, PLAUR, CD38*, and *HLA* genes, are associated with CD11c^+^CCR2^+^ and CD11c^+^CD38^+^ mass cytometry clusters (**Fig. 5h**). These markers may define inflammatory synovial monocyte populations. We further associated proteomic expression of monocytes with distinct scRNA-seq clusters based on marker genes (AUC>0.7) and observed that population defined by cell surface CD11c^+^CD38^+^ is highly associated with the activated monocytes states (SC-M1 and SC-M4) (z-score=2.3) (**Fig. 5i, Table 1**). Supporting this finding, indeed using MASC, we confirmed that synovial CD11c^+^CD38^+^ monocytes are significantly expanded in leukocyte-rich RA (OR = 7.8 (95% CI: 3.6-17.2), one tailed MASC p=6.7e-05) (**Table 1**). Conversely, monocytes from cluster SC-M2 correlate with CD11c^−^ by mass cytometry and are inversely correlated with inflammatory monocyte populations (z-score=2.7) (**Fig. 5i, Table 1**).

### Heterogeneity in synovial CD4 and CD8 T cells defined by effector functions

Single-cell RNA-seq data defined distinct CD4^+^ and CD8^+^ T cell populations (**Fig. 6a**). Among CD4^+^ T cells, expression of *CCR7* and *SELL* were notably higher in SC-T1 and the central memory T cells gene set is enriched in SC-T1 expressed genes (**Fig. 6a,b**), supporting the identification of SC-T1 as central memory T cells. Next, we identified two populations of CD4^+^ T cells marked by high expression of *FOXP3* (SC-T2) and *CXCL13* (SC-T3) (**Fig. 6b**). Examination of differentially expressed genes between these two T cell subsets suggested that SC-T2 represents *FOXP3^+^* Tregs^33^ while SC-T3 represents *PD-1^+^* Tph cells and Tfh cells^20^ (**Supplemental Fig. 6**).

**Fig. 6.**
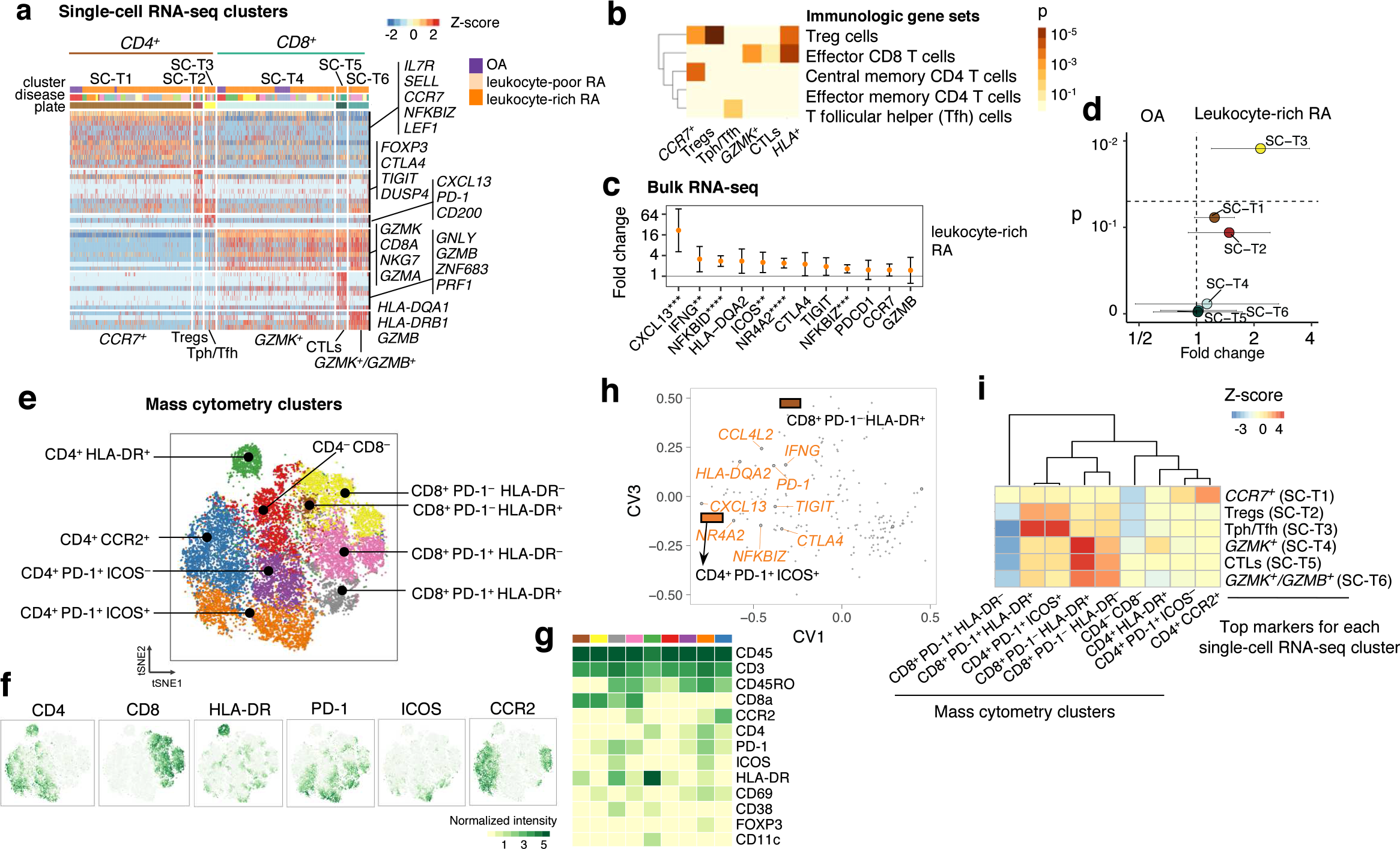
Synovial T cells display heterogeneous subpopulations in RA synovium, **a.** Single-cell RNA-seq analysis identified three *CD4^+^* subsets: *CCR7^+^* (SC-T1), Treg (SC-T2), and Tph/Tfh (SC-T3); and three *CDS^+^* subsets: *GZMK\** (SC-T4), CTLs (SC-T5), and *GZMK^+^/GZMB^+^* (SC-T6). **b.** Pathway analysis based on immunologic gene set enrichment indicates the potential enriched T cell states pathways. The brief description of the standard names from up to bottom are: Genes up-regulated in CD4 high cells from thymus: Treg versus T conv (GSE42021); Genes up-regulated in comparison of effector CD8 T cells versus memory CD8 T cells (GOLDRATH); Genes down-regulated in comparison of effector memory T cells versus central memory T cells from peripheral blood mononuclear cells (PBMC) (GSE11057); Genes up-regulated in comparison of effective memory CD4 T cells versus Th1 cells (GSE3982); Genes up-regulated in comparison of T follicular helper (Tfh) cells versus Th17 cells (GSE11924). **c.** Differential expression analysis on leukocyte-rich RA (n=18) comparing with OA (n=13) on sorted T cell samples revealed that *CXCL13, NFKBID, NFKBIZ*, and *NR4A2* are significantly upregulated in leukocyte-rich RA. ****p ≤ 1e-4, ***p ≤ 1e-3, **p ≤ 1e-2. **d.** Disease association of single-cell RNA-seq clusters by aggregating top markers (AUC>0.7) by comparing leukocyte-rich RA with OA using bulk RNA-seq. Tph/Tfh cells (SC-T4) are upregulated (two-tailed Student’s *t*-test p=0.01, t-value=2.73, df =29.00) in leukocyte-rich RA. **e.** Mass cytometry analysis by DensVM revealed nine T cell subpopulations, **f-g.** Distinct pattern of protein markers that used to define these clusters, **h.** Integration of identified mass cytometry clusters with bulk RNA-seq using CCA reveals bulk genes that are associated with CD4^+^PD-1^+^ ICOS^+^ and CD8^+^ PD-1^−^ HLA-DR^+^ by mass cytometry, **i.** Integration of mass cytometry clusters with single-cell RNA-seq clusters by taking the average of the top markers (AUC>0.7) for each single-cell RNA-seq cluster in the top 10 canonical variates. Z-score based on permutation test reveals that CD4^+^ PD-1^+^ICOS^+^ and CD8^+^ PD-1^+^ HLA-DR^+^ by mass cytometry are highly associated with Tph/Tfh (SC-T3) by single-cell RNA-seq; CD8^+^PD-1^−^ HLA-DR^+^ T cells by mass cytometry are highly associated with CD8^+^ T cells (SC-T4, SC-T5, and SC-T6).

Single-cell RNA-seq analysis of synovial CD8^+^ T cells identified three unexpected populations characterized by distinct expression of effector molecules *GZMK, GZMB, GZMA* and *GNLY* (**Fig. 6a**). We defined these populations as *GZMK^+^CD8^+^* (SC-T4), *GNLY^+^GZMB^+^* cytotoxic T lymphocytes (CTLs) (SC-T5), and *GZMK^+^/GZMB^+^* T cells (SC-T6). *GZMK^+^/GZMB^+^* T cells not only expressed *HLA-DRA* and *HLA-DQA1* at high levels, but also expressed genes suggestive of an effector phenotype (**Fig. 6a,b**). Application of GSEA to these populations annotated each of the six T cell clusters (**Fig. 6b**).

Many genes specifically expressed by T cell subsets in bulk-sorted T cells were upregulated in leukocyte-rich RA synovia comparing to OA by bulk-sorted T cells (CD45^+^CD14^−^ CD3^+^), including chemokine *CXCL13* (*t*-test p=1.2e-4) and *NFKBID* (*t*-test p=1.6e-6), a gene downstream of TCR activation (**Fig. 6c**). This likely reflected expansion of Tph cells and activated T cell subsets. Indeed, unbiased interrogation of bulk RNA-seq T cell data using the top differentially expressed genes (AUC>0.7) among scRNA-seq T cell subsets revealed significant expansion of Tph/Tfh cells (*t*-test p=0.01) (**Fig. 6d**).

Using mass cytometry, we identified nine putative T cell clusters among the synovial T cells (CD45^+^CD14^−^CD3^+^) (**Fig. 6e-g, Supplemental Fig. 4c**). By integrating bulk RNA-seq with mass cytometry cluster abundances, we found that the gene expression of *CXCL13* and inhibitory receptors *TIGIT*and *CTLA4* are associated with abundance of the CD4^+^PD-1^+^ICOS^+^ mass cytometry cluster. The abundance of CD8^+^HLA-DR^+^ cells was associated with the expression of gene *IFNG* and *HLA-DQA2* (**Fig. 6h**). When aggregating the differentially expressed marker genes (AUC>0.7) for scRNA-seq clusters, we consistently observed significant associations between Tph/Tfh cells (SC-T3) and CD4^+^PD-1^+^ICOS^+^ T cells (z-score = 3.4); CD8^+^ subsets including *GZMK^+^/GZMB^+^* (SC-T6), CTLs (SC-T5), and *GZMK^+^* (SC-T4) and CD8^+^PD-1^−^HLA-DR^+^ T cells by mass cytometry (**Fig. 6i, Table 1**), confirming their respective identities. In addition, CD4^+^PD-1^+^ICOS^+^ cells were significantly expanded in leukocyte-rich RA (MASC OR = 3 (95% CI: 1.7-5.2), one tailed MASC p=2.7e-04) (**Table 1**). Interestingly, Tregs (SC-T2) exhibited nominal association with CD4^+^PD-1^+^ICOS^+^ and CD8^+^PD1^+^HLA-DR^+^ T cells (z-score = 1.5), potentially due to shared gene expression programs between Tregs, Tph/Tfh, and CD8^+^PD-1^+^HLA-DR^+^ T cells.

### Autoimmune-associated B cells expanded in RA synovium by single-cell RNA-seq

We identified synovial B cell 4 clusters with scRNA-seq: naïve B cells (SC-B1), memory B cells (SC-B2), *CD11c^+^* ABC cells (SC-B3), and plasmablasts (SC-B4) (**Fig. 7a**). Using Gene Ontology pathway enrichment for these four subsets we observed that MHC Class II protein complex and interferon-gamma-mediated signaling pathways (**Supplemental Fig. 5b**) were enriched in all the *HLA^+^* subsets, SC-B1, SC-B2, and SC-B3 (**Fig. 3f**), suggesting B cell activation. Pathway analysis on curated immunological genes sets demonstrated that SC-B1 expresses naïve B cell genes, while SC-B2 and SC-B3 are more enriched for IgM and IgG memory B cell genes (**Fig. 7b**). Intriguingly, we observed that SC-B3 cells express high levels of *CD11c* and *T-bet* (**Fig. 7b**), which are autoimmune-associated B cells (ABC) markers^30,31^, as well as markers of recently activated B cells including ACTB^34^. High expression of *AICD* is also in accord with the recently reported transcriptomic analysis of CD11c^+^ B cells from SLE peripheral blood^35^. While ABCs constitute as a relatively small proportion of all B cells, they are almost exclusively derived from two leukocyte-rich RA patients. Examination of bulk transcriptomic profiles of synovial B cell samples shows that genes *MZB1, XBP1* and *CD11c* genes are upregulated in leukocyte-rich RA (n=16) compared to OA (n=7) (**Fig. 7c**).

**Fig. 7.**
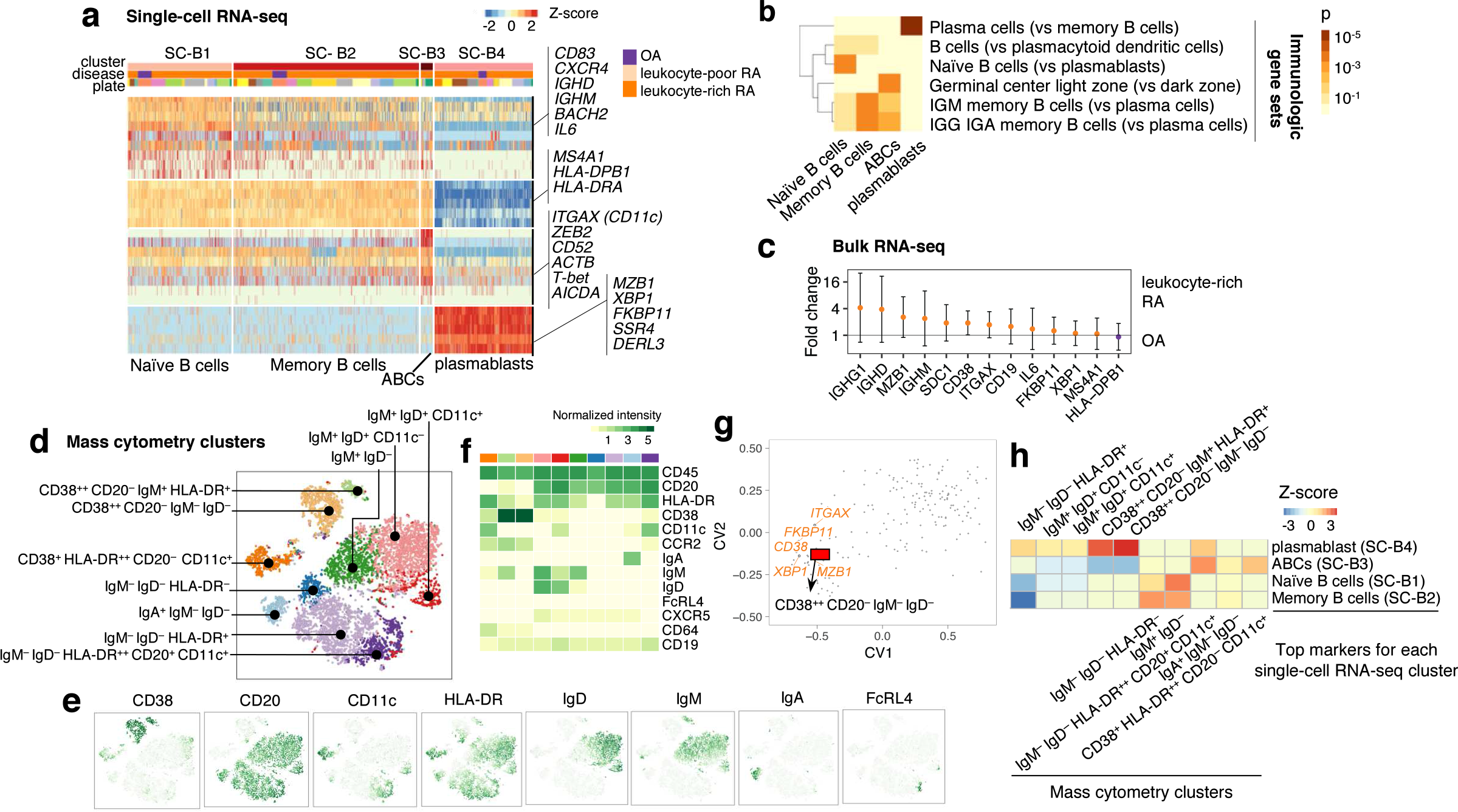
Synovial B cells display heterogeneous subpopulations in RA synovium, **a.** Single-cell RNA-seq analysis identified naive B cells (SC-B1), memory B cells (SC-B2), autoimmune-associated B cells (ABCs) (SC-B3), and plasmablasts (SC-B4). **b.** Pathway enrichment analysis using immunologic gene sets indicates the distinct enriched pathways for each single-cell RNA-seq cluster. The standard names for the immunological gene sets from up to bottom are: Genes up-regulated in plasma cells versus memory B cells (GSE12366); Genes up-regulated in comparison of B cells versus plasmacytoid dendritic cells (pDC) (GSE29618); Genes up-regulated in B lymphocytes: naive versus plasmablasts (GSE42724); Genes up-regulated in B lymphocytes: human germinal center light zone versus dark zone (GSE38697); Genes up-regulated in comparison of memory IgM B cells versus plasma cells from bone marrow and blood (GSE22886); Genes up-regulated in comparison of memory IGG and IGA B cells versus plasma cells from bone marrow and blood (GSE22886). **c.** Differential expression analysis by comparing leukocyte-rich RA (n=16) with OA (n=7) by bulk RNA-seq. **d.** Mass cytometry data analysis identified ten clusters by DensVM. **e-f.** Distinct expression patterns of protein markers that used to define these clusters, **g.** Integrating mass cytometry clusters with bulk RNA-seq data using CCA shows that CD38^+^ CD20^−^ lg^−^ (plasmablasts) is highly associated with gene expression of plasma cells makers, like *XBP1*. **h.** Integration of mass cytometry clusters with single-cell RNA-seq clusters suggested that CD38^+^ CD20^−^ lgM^+^ and CD38^+^ CD20^−^ lg^−^ are significantly associated with plasmablast (SC-64); HLA-DR^high^ CD20^+^ CD11c^+^ B cells are associated with ABCs (SC-B3).

Mass cytometric data of synovial B cells (CD45^+^CD3^−^CD14^−^CD19^+^) identified ten putative B cell clusters (**Fig. 7d-f**, **Supplemental Fig. 4d**). Next, we analyzed bulk RNA-seq and mass cytometry cluster abundances from the shared samples, and found that the gene expressions of *CD38, MZB1*, and plasma cell differentiation factor *XBP1* are associated with abundance of CD38^++^CD20^−^IgM^−^IgD^−^ plasmablasts (**Fig. 7g**). To further validate the distinct scRNA-seq clusters using mass cytometry, we integrated ten mass cytometry populations with scRNA-seq clusters and observed significant correlation between plasmablasts (SC-B4) and CD38^++^CD20^−^IgM^−^IgD^−^ B cells (z-score=2.7) (**Fig. 7h, Table 1**). Consistent with identification of ABCs in RA synovia, *CD11c*^+^ ABCs (SC-B3) were positively correlated (z-score=1.6) with IgM^−^ IgD^−^ HLA-DR^++^ CD20^+^ CD11c^+^, which is significantly (OR = 5.7 (95% CI: 1.8-22.3), one tailed MASC p=2.7e-03) expanded in leukocyte-rich RA (**Fig. 7h**, **Table 1**). Mass cytometry analysis further identified three putative subsets within CD11c^+^ cells based on expression of immunoglobulin profiles: IgM^−^ IgD^−^ HLA-DR^++^ CD20^+^ CD11c^+^, CD38^+^ HLA-DR^++^ CD20^−^ CD11c^+^, and IgM^+^ IgD^+^ CD11c, suggesting additional heterogeneity within ABCs. Among these, IgM^+^ IgD^+^ CD11c B cells express FcRL4, suggesting homology to a population of CD11c^+^FcRL4^+^ memory B cells described in the human tonsil.

### Inflammatory pathways and effector modules revealed by global transcriptomic profiling

We used bulk and single cell transcriptomes of sorted synovial cells to detect pathogenic molecular signal pathways. First, principal component analysis (PCA) on post-QC OA and leukocyte-rich RA samples (**Supplemental Fig. 7a,b**) demonstrated that cell type accounted for most of the variance and each cell type expressed specific marker genes (**Supplemental Fig. 7c**). Within each cell-type we observed that leukocyte-rich RA appeared distinct from OA samples, but leukocyte-poor RA grouped together with OA samples (**Supplemental Fig. 7d-g**). We observed that 173 genes in fibroblasts, 159 genes in monocytes, 10 genes in T cells, and 5 genes in B cells were upregulated in leukocyte-rich RA tissues compared to OA (FC>2 and FDR<0.01). To define the pathways relevant to leukocyte-rich RA, we applied GSEA weighted by gene effect sizes and identified TLR signaling (monocytes and B cells), type I interferon response and inflammatory response (monocytes and fibroblasts) (**Supplemental Fig. 7h-i**), Fc receptor signaling (monocytes), NF-kappa B signaling (fibroblasts), and interferon gamma (T cells) pathways (**Fig. 8a**). We observed that in fibroblasts and monocytes that inflammatory response genes (*PTGS2, PTGER3*, and *ICAM1*), interferon response genes (*IFIT2, RSAD2, STAT1*, and *XAF1*), and chemokine/cytokine genes (*CCL2* and *CXCL9*) were significantly upregulated in leukocyte-rich RA (**Fig. 8b**), suggesting a coordinated chemotactic response to interferon activation. We also observed upregulation of interferon regulatory factors (IRFs), including *IRF7* and *IRF9* in T cells, and *IRF1, IRF7, IRF8* and *IRF9* in monocytes. Synovial monocytes in leukocyte-rich RA exhibit increased expression of *TLR8* and *MYD88*, consistent with IL-1 or TLR signaling (**Fig. 8a**). Taken together, pathway analysis suggests crosstalk between immune and stromal cells in leukocyte-rich RA synovia. Inflammatory response genes upregulated in leukocyte-rich RA, had comparable expression in leukocyte-poor RA and OA synovial cells (**Fig. 8b**), suggesting leukocyte infiltration is a key drive of molecular heterogeneity in RA synovia.

**Fig. 8.**
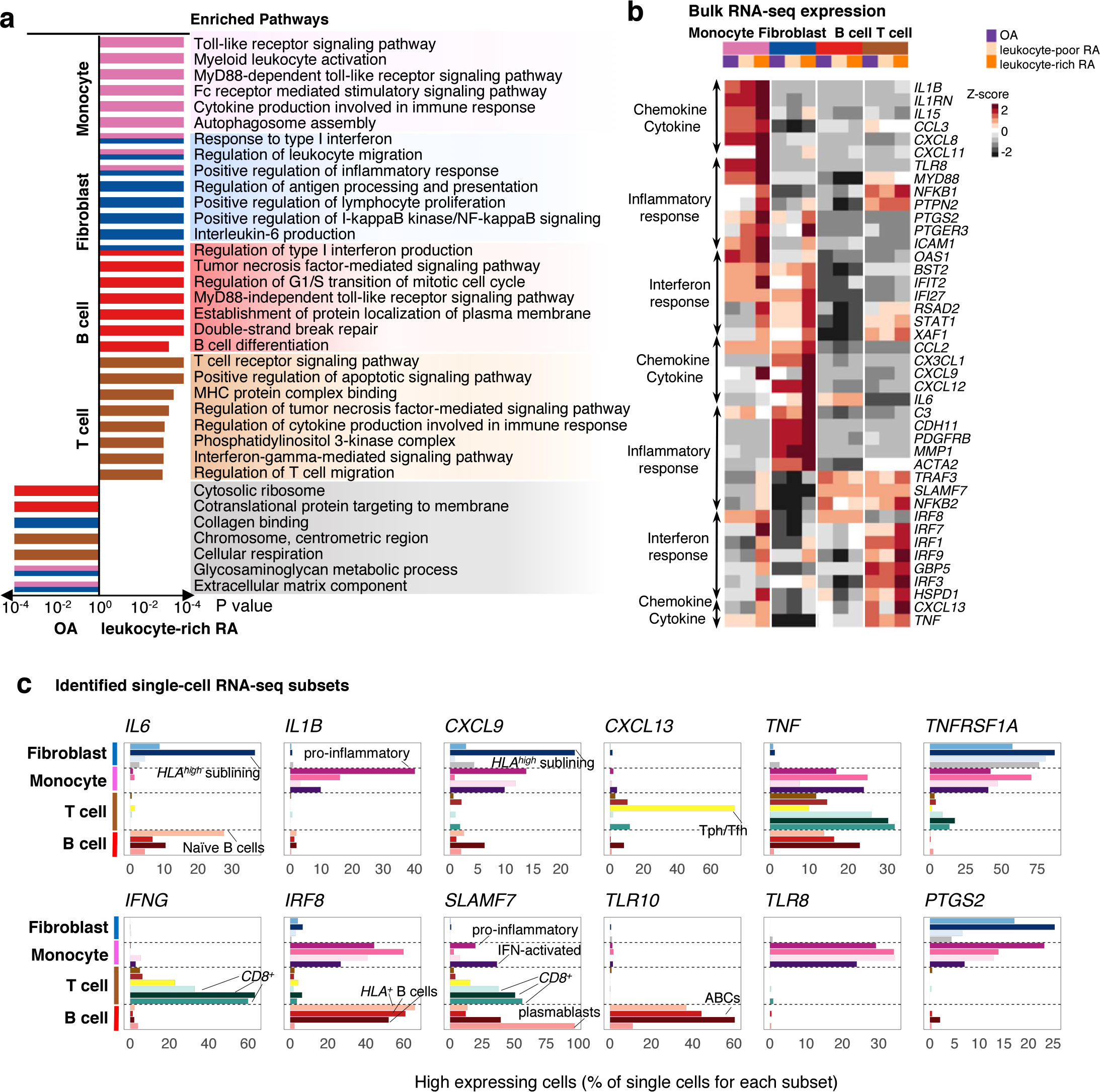
Transcriptomic profiling of synovial cells reveal upregulation of inflammatory pathways in RA synovium, **a.** Pathway enrichment driven by PCA analysis and differential expression analysis using bulk RNA-seq identified shared and unique inflammatory response pathways for each cell type. **b.** Bulk RNA-seq profiling of genes obtained from the significantly enriched pathways from (**a**) shows the averaged gene expression from different disease cohorts (OA, leukocyte-poor RA, and leukocyte-rich RA) normalized across all cell type samples, **c.** Single-cell RNA-seq profiling resolved that inflammatory cytokines/chemokines, interferon responsive, and inflammatory responsive genes were driven by a global upregulation within a synovial cell type or discrete cell states.

Next, we asked whether inflammatory cytokines upregulated in leukocyte-rich RA are driven by global upregulation within a synovial cell type, or specific upregulation within a discrete cell subset defined by scRNA-seq. Whereas *TNF* was produced at a high level by a multiple monocyte, B cell and T cell populations; *IL6* expression was restricted to *HLA^hgh^* sublining fibroblasts (SC-F2) and naive B cells (SC-B1) (**Fig. 8c**). Similarly, expression of *IL1B* and *CXCL13* was restricted to *IL1B^+^* pro-inflammatory monocytes (SC-M1) and Tph/Tfh cells (SC-T3), respectively. Surprisingly, we identified CD8 T cells, rather than CD4 T cells, as the dominant source of *IFNγ* in leukocyte-rich synovia.

We also observed cell subset-specific responses to inflammatory pathways. Toll-like receptor signaling pathway was strongly enriched in B cells and monocytes in leukocyte-rich RA tissues (**Fig. 8a**). At the single cell level, we observed that *TLR10* was only expressed by *HLA^+^* B cells, indicating that *TLR10* has a functional role within the B cell lineage^36^. In contrast, *TLR8* was significantly elevated in all RA monocyte subsets. The hematopoietic cell-specific transcription factor *IRF8* was expressed in a significant fraction of monocytes and *HLA^+^* B cells that cooperatively regulate differentiation of monocytes and activated B cells in RA synovium. *SLAMF7*, a potential therapeutic target for Systemic Lupus Erythematosus (SLE)^37^, is highly expressed by pro-inflammatory monocytes (SC-M1), IFN-activated monocytes (SC-M4), plasmablasts (SC-B4) and CD8 T cells.

## DISCUSSION

Using multi-model, high-dimensional synovial tissue data we defined stromal and immune cell populations expanded in RA indicating essential inflammatory pathways. Recognizing the considerable variation in clinical parameters for disease duration and activity, treatment types, and joint histology scores^38,39^, we elected to use a molecular parameter, based on percent lymphocytes, monocytes of the total cellularity, to classify our samples at the local tissue level. We note that differences in leukocyte enrichment of joint replacement samples and biopsy samples were best explained by leukocyte infiltration and not by the tissue source (**Supplemental Fig. 1, Supplemental Fig. 7d-g**).

This and previous studies have highlighted stromal fibroblasts as a potential therapeutic target in RA^40,12^. Consistent with previous reports^12,23,41^, we identified sublining fibroblasts as a major producer of pro-inflammatory cytokines, notably *IL6*, within the leukocyte-rich synovium (**Fig. 4**). Furthermore, a single subset of those fibroblasts expressing MHC II (SC-F2, THY1^+^CD34^−^HLA-DR^high^) was >15 fold expanded in RA tissues, highlighting it as a possible therapeutic target. In addition, MHC II expression supports a role for stromal cells in T cell antigen presentation^42^. We also observed that T cells, B cells, and monocyte proportions track with synovial fibroblasts gene expression, suggesting that synovial fibroblasts respond to infiltrating lymphocytes in RA synovium (**Supplemental Fig. 8**). Intriguingly, *DNASE1L3*, a gene whose loss of function is associated with RA^43^ and systemic lupus erythematosus^44^ susceptibility in recent genetic studies, was found to be highly expressed in synovial *CD55^+^* lining fibroblasts (SC-F4), which was relatively depleted in human RA. We identified a novel fibroblast subset (SC-F3) characterized by high *DKK3* (**Fig. 4**), encoding Dickkopf3, and protein upregulated in OA that prevents cartilage degradation in vitro^45,46^.

Transcriptional heterogeneity in the synovial monocyte compartment indicated that distinct RA-enriched subsets are driven by inflammatory cytokines (such as IL-1 or TNF) and interferons (**Fig. 5, Fig. 8**). This suggests monocyte may be sensitive to the local microenvironment with unique cytokine combinations constituting the inflammatory milieu in the RA synovium. These inflammatory phenotypes align with effective RA therapeutic targets, for example TNF and the interferon-activated JAK kinases, respectively^47,48^. The *NUPR1^+^* monocytes demonstrated lower proportions in RA tissue and had transcriptomes that were anticorrelated with the inflammatory phenotypes, suggesting either an anti-inflammatory phenotype, supported by high levels of *MERTK* (**Fig. 5**)^49^, or an unrecognized monocyte phenotype specific to the normal uninflamed synovium. Alternatively, *NUPR1^+^* markers such as osteoactivin (*GPNMB*) and cathepsin K (*CTSK*) could indicate a specific subset of osteoclast progenitors that control bone remodeling (**Fig. 5**)^50,48,51^. Further studies on normal and various disease control synovial tissues may clarify the functional role of the *NUPR1^+^* (SC-M2) monocyte phenotype. Furthermore, anatomical and spatial studies of the identified monocyte populations—particularly focused on lining versus sublining, perivascular and lymphocyte aggregate-associated monocytes—will help to elevate our understanding of the functional roles for these myeloid cell types.

Single cell classification of T cell subsets in RA synovium demonstrated CD4^+^ T cell heterogeneity that is consistent with distinction between the homing capacity and effector functions of these subsets. Consistent with previous studies, we observed expansion of CD4^+^ T peripheral helper cells^20^ (SC-T3, *CD4^+^PD-1^+^ICOS^+^*) within leukocyte-rich RA synovium. We also identified distinct CD8 T cell subsets (SC-T4-6) characterized by high expression of *IFNG* and a distinct granzyme expression pattern (**Fig. 6**). A larger study may be better powered to differentiate the relative expansion of individual subpopulations. A role of CD8^+^ T cells is consistent with MHC class I genetic associations in rheumatoid arthritis^52^, and may be relevant to tissue inflammation.

To our knowledge, this study is the first to report the presence of autoimmune-associated B cells (SC-B3, ABCs) by transcriptomic sequencing data in leukocyte-rich synovium in RA. This B cells population, dependent on *T-bet*for generation and expressing *CD11c*, was first reported in aging mice; subsequently it was seen to be expanded in autoimmune mice and enriched for autoreactive specificities^53,54^. We observed heterogeneity in this cell subset, with a sizable population of *CD11c^+^* B cells detectable in both *IgD^+^* and switched B cell populations by mass cytometry. The expression of other markers by ABCs in our transcriptome analysis suggests a balance between germinal center (*IRF8, AID*)^55^ and plasma cell (*SLAMF7*) differentiation programs within the RA synovium. We observed that multiple B cell subsets expressed MHC II, consistent with the potential for B cell antigen presentation in the RA target tissue. As previously reported, we observed in leukocyte-rich RA synovium an expansion of plasma cells^56^ (SC-B4), which are targeted by rituximab^57^, an effective RA therapy, as previously demonstrated. We also observe that naive B cells are a dominant *IL6* producer. In contrast to leukocyte-rich RA, OA synovia contain comparatively few B cells (**Fig. 2b**), which limited our ability to identify RA-associated synovial B cell subsets through case-control comparisons (**Fig. 7g**).

A critical unmet need in RA is identifying therapeutic targets for patients failing to respond to DMARDs and biologics^38^. We observed upregulation of chemokines (*CXCL8*, *CXCL9*, and *CXCL13*), cytokines (*IFNG* and *IL15*), and surface receptors (*PDGFRB* and *SLAMF7*) in distinct immune and stromal cell populations, suggesting potential novel targets. This study was enabled by important advances in the statistical analysis of single-cell data^21,58–61^ alongside rapid improvements in scaling single cell technologies^17,62^ and our recent work optimizing robust methodologies for disaggregation of synovial tissue^24^.

We advance strategies to integrate multiple molecular data sets; these approaches modulate the effect of technical artifact, frequently confounding single cell technologies^63–65^, while emphasizing biological signals. Our CCA-based integrative strategy clusters highdimensional scRNA-seq data using canonical variates that capture variance that are present in both the single-cell and bulk RNA-seq data. These shared variances likely represent biological trends, and not technical factors that would likely be uncorrelated in these two independent data sets. CCA has been successfully employed effectively in other contexts to integrate highdimensional biological data^66,65^. Penalized CCA^67^ and deep CCA^68,69^ can produce non-linear variates and may prove to be highly effective as we confront higher throughput platforms with greater cell-to-cell data.

The two single cell modalities used in this study, mass cytometry and scRNA-seq, complement each other. Single-cell RNA-seq captures expression of thousands of genes^70,71^, but at the cost of sparse data^63^. A single mass cytometry assay captures hundreds of thousands of individual cells, but only measures a limited number (~40)^72^ of pre-selected markers. But, since markers are backed with decades of experimental experience they can be effective at defining cellular heterogeneity^73^. Mass cytometry analysis across all cell populations identified that leukocyte-rich patients show high cell abundances of HLA-DR^+^ fibroblast populations, Tph cells, CD11c^+^CD14^+^ monocytes, and CD11c^+^ B cell populations (**Supplemental Fig. 4e**). Combining mass cytometry with the extended dimensionality of scRNA-seq analyses, enables quantification of well-established cell populations, while also enabling discovery of rare or novel cell states, such as the CD8 T cell states noted here. We note the recent development of approaches to capture mRNA and protein expression simultaneously that will further augment our ability query tissue inflammation^74,75^.

Whether cell population expansions and molecular pathways highlighted in this study represent RA pathogenesis or a downstream effect of inflammation warrants further investigation. The RA/SLE AMP is now engaged in obtaining a large collection of synovial biopsy specimens and paired blood samples from 150 RA patients for single cell analyses with detailed clinical data, disease activity metrics, and ultrasound score evaluation of synovitis. We anticipate that this ongoing larger study will enable us to not only define additional subpopulations, but to better define their link to clinical sub-phenotypes.

It is essential to interrogate the tissue infiltration of diseases other than RA, including systemic lupus erythematosus, type I diabetes, psoriasis, multiple sclerosis and other organ targeting conditions. Application of multiple single cell technologies together can help to define key novel populations, thereby providing new insights about etiology and potential therapies.

## ACKNOWLEDGMENTS

This work was supported by the Accelerating Medicines Partnership (AMP) in Rheumatoid Arthritis and Lupus Network. AMP is a public-private partnership (AbbVie Inc., Arthritis Foundation, Bristol-Myers Squibb Company, Lupus Foundation of America, Lupus Research Alliance, Merck Sharp & Dohme Corp., National Institute of Allergy and Infectious Diseases, National Institute of Arthritis and Musculoskeletal and Skin Diseases, Pfizer Inc., Rheumatology Research Foundation, Sanofi and Takeda Pharmaceuticals International, Inc.) created to develop new ways of identifying and validating promising biological targets for diagnostics and drug development Funding was provided through grants from the National Institutes of Health (UH2-AR067676, UH2-AR067677, UH2-AR067679, UH2-AR067681, UH2-AR067685, UH2-AR067688, UH2-AR067689, UH2-AR067690, UH2-AR067691, UH2-AR067694, and UM2-AR067678). This work is also supported in part by funding from the Ruth L. Kirschstein National Research Service Award (F31AR070582) from the National Institute of Arthritis and Musculoskeletal and Skin Diseases (K.S.). K.W. is supported by a Rheumatology Research Foundation Scientist Development Award. D.A.R. is supported by NIAMS K08 AR072791-01. L.T.D. is supported by NIAMS K01 AR066063. J.H.A. is supported by R21 AR071670, and the Bertha and Louis Weinstein research fund. K.S. is supported by NIAMS F31-AR070582. S.R. is supported by 1R01AR063759-01A1 and Doris Duke Charitable Foundation Grant #2013097. A.F., C.D.B. and J.D.T. were supported by the Arthritis Research UK Rheumatoid Arthritis (#20298), and by the National Institute for Health Research (NIHR)’s Birmingham Biomedical Research Centre program, supported by the National Institute for Health Research/Wellcome Trust Clinical Research Facility at University Hospitals Birmingham NHS Foundation Trust.

## AUTHOR CONTRIBUTIONS

S.K., S.G., D.T., L.B.H., K.S.-E., A.M.M., D.L.B., J.H.A., V.P.B., M.H., A.F., C.P., H.P., G.S.F., L.M., P.K.G., W.A. and L.T.D. recruited patients and obtained synovial tissues. B.B., E.D. and E.G. performed histological assessment of tissues. K.W., D.A.R., G.W., and M.B.B. designed and implemented tissue processing and cell sorting pipeline. J.A.L. obtained mass cytometry data from samples. N.H. and C.N. obtained single cell RNA-seq data from samples. F.Z., K.S., C.Y.F., D.J.L. and S.R. conducted computational and statistical analysis. K.S. implemented the website. S.R., M.B.B., J.H.A., and L.T.D. supervised the research. F.Z., K.W., and S.R. wrote the initial draft; K.S, C.Y.F. D.A.R, L.T.D., J.H.A, M.B.B. edited it, and all the authors participated in writing the final manuscript.

## References

1. Gibofsky, A. Epidemiology, pathophysiology, and diagnosis of rheumatoid arthritis: A Synopsis. Am. J. Manag. Care 20, 128–35 (2014).

2. McInnes, I. B. & Schett, G. The pathogenesis of rheumatoid arthritis. N. Engl. J. Med. 365, 2205–2219 (2011).

3. Orr, C. et al. Synovial tissue research: a state-of-the-art review. Nat. Rev. Rheumatol. 13, 463–475 (2017).

4. Wolfe, F. et al. The mortality of rheumatoid arthritis. Arthritis Rheum. 37, 481–494 (1994).

5. Namekawa, T., Wagner, U. G., Goronzy, J. J. & Weyand, C. M. Functional subsets of CD4 T cells in rheumatoid synovitis. Arthritis Rheum. 41, 2108–2116 (1998).

6. Gizinski, A. M. & Fox, D. A. T cell subsets and their role in the pathogenesis of rheumatic disease. Curr. Opin. Rheumatol. 26, 204–210 (2014).

7. Reparon-Schuijt, C. C. et al. Secretion of anti-citrulline-containing peptide antibody by B lymphocytes in rheumatoid arthritis. Arthritis Rheum. 44, 41–47 (2001).

8. Mulherin, D., Fitzgerald, O. & Bresnihan, B. Synovial tissue macrophage populations and articular damage in rheumatoid arthritis. Arthritis Rheum. 39, 115–124 (1996).

9. Kinne, R. W., Bräuer, R., Stuhlmüller, B., Palombo-Kinne, E. & Burmester, G. R. Macrophages in rheumatoid arthritis. Arthritis Res. 2, 189–202 (2000).

10. Müller-Ladner, U. et al. Synovial fibroblasts of patients with rheumatoid arthritis attach to and invade normal human cartilage when engrafted into SCID mice. Am. J. Pathol. 149, 1607–1615 (1996).

11. Pap, T., Müller-Ladner, U., Gay, R. E. & Gay, S. Fibroblast biology. Role of synovial fibroblasts in the pathogenesis of rheumatoid arthritis. Arthritis Res. 2, 361–367 (2000).

12. Noss, E. H. & Brenner, M. B. The role and therapeutic implications of fibroblast-like synoviocytes in inflammation and cartilage erosion in rheumatoid arthritis. Immunol. Rev. 223, 252–270 (2008).

13. Dennis, G., Jr et al. Synovial phenotypes in rheumatoid arthritis correlate with response to biologic therapeutics. Arthritis Res. Ther. 16, R90 (2014).

14. Orange, D. E. et al. Machine learning integration of rheumatoid arthritis synovial histology and RNAseq data identifies three disease subtypes. Arthritis Rheumatol (2018). doi:10.1002/art.40428

15. Lindberg, J. et al. Variability in synovial inflammation in rheumatoid arthritis investigated by microarray technology. Arthritis Res. Ther. 8, R47 (2006).

16. Stephenson, W. et al. Single-cell RNA-seq of rheumatoid arthritis synovial tissue using low-cost microfluidic instrumentation. Nat. Commun. 9, 791 (2018).

17. Papalexi, E. & Satija, R. Single-cell RNA sequencing to explore immune cell heterogeneity. Nat. Rev. Immunol. 18, 35–45 (2018).

18. Schelker, M. et al. Estimation of immune cell content in tumour tissue using single-cell RNA-seq data. Nat. Commun. 8, 2032 (2017).

19. Wong, M. T. et al. A High-Dimensional Atlas of Human T Cell Diversity Reveals Tissue-Specific Trafficking and Cytokine Signatures. Immunity 45, 442–456 (2016).

20. Rao, D. A. et al. Pathologically expanded peripheral T helper cell subset drives B cells in rheumatoid arthritis. Nature 542, 110–114 (2017).

21. Fonseka, C. Y. et al. Reverse Association Of Single Cells To Rheumatoid Arthritis Accounting For Mixed Effects Identifies An Expanded CD27-HLA-DR^+^ Effector Memory CD4^+^ T Cell Population. bioRxiv 172403 (2017). doi:10.1101/172403

22. Villani, A.-C. et al. Single-cell RNA-seq reveals new types of human blood dendritic cells, monocytes, and progenitors. Science 356, (2017).

23. Mizoguchi, F. et al. Functionally distinct disease-associated fibroblast subsets in rheumatoid arthritis. Nat. Commun. 9, 789 (2018).

24. Donlin, L. T. et al. High dimensional analyses of cells dissociated from cryopreserved synovial tissue. bioRxiv 284844 (2018). doi:10.1101/284844

25. Becher, B. et al. High-dimensional analysis of the murine myeloid cell system. Nat. Immunol. 15, 1181–1189 (2014).

26. van Baarsen, L. G. M. et al. Synovial tissue heterogeneity in rheumatoid arthritis in relation to disease activity and biomarkers in peripheral blood. Arthritis Rheum. 62, 1602–1607 (2010).

27. De Maesschalck, R., Jouan-Rimbaud, D. & Massart, D. L. The Mahalanobis distance. Chemometrics Intellig. Lab. Syst. 50, 1–18 (2000).

28. Krenn, V. et al. Grading of Chronic Synovitis — A Histopathological Grading System for Molecular and Diagnostic Pathology. Pathology - Research and Practice 198, 317–325 (2002).

29. Maaten, L. van der & Hinton, G. Visualizing Data using t-SNE. J. Mach. Learn. Res. 9, 2579–2605 (2008).

30. Rubtsov, A. V. et al. CD11c-Expressing B Cells Are Located at the T Cell/B Cell Border in Spleen and Are Potent APCs. J. Immunol. 195, 71–79 (2015).

31. Pillai, S. Now you know your ABCs. Blood 118, 1187–1188 (2011).

32. Todd, D. J. et al. XBP1 governs late events in plasma cell differentiation and is not required for antigen-specific memory B cell development. J. Exp. Med. 206, 2151–2159 (2009).

33. Josefowicz, S. Z., Lu, L.-F. & Rudensky, A. Y. Regulatory T cells: mechanisms of differentiation and function. Annu. Rev. Immunol. 30, 531–564 (2012).

34. Ellebedy, A. H. et al. Defining antigen-specific plasmablast and memory B cell subsets in human blood after viral infection or vaccination. Nat. Immunol. 17, 1226–1234 (2016).

35. Wang, S. et al. IL-21 drives expansion and plasma cell differentiation of autoreactive CD11c hi T-bet^+^ B cells in SLE. Nat. Commun. 9, 1758 (2018).

36. Hess, N. J., Jiang, S., Li, X., Guan, Y. & Tapping, R. I. TLR10 Is a B Cell Intrinsic Suppressor of Adaptive Immune Responses. J. Immunol. 198, 699–707 (2017).

37. Comte, D. et al. Signaling Lymphocytic Activation Molecule Family Member 7 Engagement Restores Defective Effector CD8^+^ T Cell Function in Systemic Lupus Erythematosus. Arthritis Rheumatol 69, 1035–1044 (2017).

38. Smolen, J. S. How well can we compare different biologic agents for RA? Nat. Rev. Rheumatol. 6, 247–248 (2010).

39. Pitzalis, C., Kelly, S. & Humby, F. New learnings on the pathophysiology of RA from synovial biopsies. Curr. Opin. Rheumatol. 25, 334–344 (2013).

40. Filer, A. The fibroblast as a therapeutic target in rheumatoid arthritis. Curr. Opin. Pharmacol. 13, 413–419 (2013).

41. Nguyen, H. N. et al. Autocrine Loop Involving IL-6 Family Member LIF, LIF Receptor, and STAT4 Drives Sustained Fibroblast Production of Inflammatory Mediators. Immunity 46, 220–232 (2017).

42. Tran, C. N. et al. Presentation of arthritogenic peptide to antigen-specific T cells by fibroblast-like synoviocytes. Arthritis Rheum. 56, 1497–1506 (2007).

43. Westra, H.-J. et al. Fine-mapping identifies causal variants for RA and T1D in DNASE1L3, SIRPG, MEG3, TNFAIP3 and CD28/CTLA4 loci. bioRxiv 151423 (2017). doi:10.1101/151423

44. Al-Mayouf, S. M. et al. Loss-of-function variant in DNASE1L3 causes a familial form of systemic lupus erythematosus. Nat. Genet. 43, 1186–1188 (2011).

45. Snelling, S. J. B. et al. Dickkopf-3 is upregulated in osteoarthritis and has a chondroprotective role. Osteoarthritis Cartilage 24, 883–891 (2016).

46. Snelling, S., Davidson, R., Swingler, T., Price, A. & Clark, I. Dkk3 Can prevent cartilage degradation and modulate TGFbeta and Wnt signalling. Osteoarthritis Cartilage 20, S10 (2012).

47. Weinblatt, M. E. et al. A trial of etanercept, a recombinant tumor necrosis factor receptor:Fc fusion protein, in patients with rheumatoid arthritis receiving methotrexate. N. Engl. J. Med. 340, 253–259 (1999).

48. Lee, E. B. et al. Tofacitinib versus methotrexate in rheumatoid arthritis. N. Engl. J. Med. 370, 2377–2386 (2014).

49. Zizzo, G., Hilliard, B. A., Monestier, M. & Cohen, P. L. Efficient clearance of early apoptotic cells by human macrophages requires M2c polarization and MerTK induction. J. Immunol. 189, 3508–3520 (2012).

50. Skoumal, M. et al. Serum cathepsin K levels of patients with longstanding rheumatoid arthritis: correlation with radiological destruction. Arthritis Res. Ther. 7, R65–70 (2005).

51. Frara, N. et al. Transgenic Expression of Osteoactivin/gpnmb Enhances Bone Formation In Vivo and Osteoprogenitor Differentiation Ex Vivo. J. Cell. Physiol. 231, 72–83 (2016).

52. Raychaudhuri, S. et al. Five amino acids in three HLA proteins explain most of the association between MHC and seropositive rheumatoid arthritis. Nat. Genet. 44, 291 (2012).

53. Rubtsov, A. V. et al. Toll-like receptor 7 (TLR7)-driven accumulation of a novel CD11c^+^ B-cell population is important for the development of autoimmunity. Blood 118, 1305–1315 (2011).

54. Manni, M. et al. Regulation of age-associated B cells by IRF5 in systemic autoimmunity. Nat. Immunol. 19, 407–419 (2018).

55. Cattoretti, G. et al. Stages of germinal center transit are defined by B cell transcription factor coexpression and relative abundance. J. Immunol. 177, 6930–6939 (2006).

56. Humby, F. et al. Ectopic Lymphoid Structures Support Ongoing Production of Class-Switched Autoantibodies in Rheumatoid Synovium. PLoS Med. 6, e1 (2009).

57. Marston, B., Palanichamy, A. & Anolik, J. H. B cells in the pathogenesis and treatment of rheumatoid arthritis. Curr. Opin. Rheumatol. 22, 307–315 (2010).

58. Dey, K. K., Hsiao, C. J. & Stephens, M. Visualizing the structure of RNA-seq expression data using grade of membership models. PLoS Genet. 13, e1006599 (2017).

59. Kiselev, V. Y. et al. SC3: consensus clustering of single-cell RNA-seq data. Nat. Methods (2017). doi:10.1038/nmeth.4236

60. Satija, R., Farrell, J. A., Gennert, D., Schier, A. F. & Regev, A. Spatial reconstruction of single-cell gene expression data. Nat. Biotechnol. 33, 495–502 (2015).

61. Wang, B., Zhu, J., Pierson, E., Ramazzotti, D. & Batzoglou, S. Visualization and analysis of single-cell RNA-seq data by kernel-based similarity learning. Nat. Methods (2017). doi:10.1038/nmeth.4207

62. Svensson, V., Vento-Tormo, R. & Teichmann, S. A. Exponential scaling of single-cell RNA-seq in the last decade. arXiv [q-bio.GN] (2017).

63. Hicks, S. C., Townes, F. W., Teng, M. & Irizarry, R. A. Missing data and technical variability in single-cell RNA-sequencing experiments. Biostatistics (2017). doi:10.1093/biostatistics/kxx053

64. Tung, P.-Y. et al. Batch effects and the effective design of single-cell gene expression studies. Sci. Rep. 7, 39921 (2017).

65. Butler, A., Hoffman, P., Smibert, P., Papalexi, E. & Satija, R. Integrating single-cell transcriptomic data across different conditions, technologies, and species. Nat. Biotechnol. (2018). doi:10.1038/nbt.4096

66. Parkhomenko, E., Tritchler, D. & Beyene, J. Sparse canonical correlation analysis with application to genomic data integration. Stat. Appl. Genet. Mol. Biol. 8, Article 1 (2009).

67. Witten, D. M., Tibshirani, R. & Hastie, T. A penalized matrix decomposition, with applications to sparse principal components and canonical correlation analysis. Biostatistics 10, 515–534 (2009).

68. Andrew, G., Arora, R., Bilmes, J. & Livescu, K. Deep Canonical Correlation Analysis. in International Conference on Machine Learning 1247–1255 (2013).

69. Benton, A. et al. Deep Generalized Canonical Correlation Analysis. arXiv [cs.LG] (2017).

70. Hashimshony, T. et al. CEL-Seq2: sensitive highly-multiplexed single-cell RNA-Seq. Genome Biol. 17, 77 (2016).

71. Zheng, G. X. Y. et al. Massively parallel digital transcriptional profiling of single cells. Nat. Commun. 8, 14049 (2017).

72. Bendall, S. C. et al. Single-cell mass cytometry of differential immune and drug responses across a human hematopoietic continuum. Science 332, 687–696 (2011).

73. Bjornson, Z. B., Nolan, G. P. & Fantl, W. J. Single-cell mass cytometry for analysis of immune system functional states. Curr. Opin. Immunol. 25, 484–494 (2013).

74. Peterson, V. M. et al. Multiplexed quantification of proteins and transcripts in single cells. Nat. Biotechnol. 35, 936–939 (2017).

75. Stoeckius, M. et al. Simultaneous epitope and transcriptome measurement in single cells. Nat. Methods 14, 865–868 (2017).

76. Finck, R. et al. Normalization of mass cytometry data with bead standards. Cytometry A 83, 483–494 (2013).

77. González, I., Déjean, S., Martin, P. & Baccini, A. CCA: An R Package to Extend Canonical Correlation Analysis. Journal of Statistical Software, Articles 23, 1–14 (2008).

78. Sing, T., Sander, O., Beerenwinkel, N. & Lengauer, T. ROCR: visualizing classifier performance in R. Bioinformatics 21, 3940–3941 (2005).

79. Subramanian, A. et al. Gene set enrichment analysis: A knowledge-based approach for interpreting genome-wide expression profiles. Proceedings of the National Academy of Sciences 102, 15545–15550 (2005).

80. Ashburner, M. et al. Gene ontology: tool for the unification of biology. The Gene Ontology Consortium. Nat. Genet. 25, 25–29 (2000).

81. Liberzon, A. et al. The Molecular Signatures Database (MSigDB) hallmark gene set collection. Cell Syst 1, 417–425 (2015).

82. Reynolds, A. P., Richards, G., de la Iglesia, B. & Rayward-Smith, V. J. Clustering Rules: A Comparison of Partitioning and Hierarchical Clustering Algorithms. J. Math. Model. Algorithms 5, 475–504 (2006).

83. Rousseeuw, P. J. Silhouettes: A graphical aid to the interpretation and validation of cluster analysis. J. Comput. Appl. Math. 20, 53–65 (1987).

